# EphrinA5 regulates cell motility by modulating the targeting of DNMT1 to the *Ncam1* promoter via lncRNA/DNA triplex formation

**DOI:** 10.1101/2023.03.25.534129

**Authors:** Can Bora Yildiz, Tathagata Kundu, Julia Gehrmann, Jannis Koesling, Amin Ravaei, Mira Jakovcevski, Daniel Pensold, Olav Zimmermann, Giulia Rossetti, Ivan G. Costa, Geraldine Zimmer-Bensch

## Abstract

Cell-cell communication is mediated by membrane receptors and their cognate ligands, such as the Eph/ephrin system, and dictates physiological processes, including cell proliferation and migration. However, whether and how Eph/ephrin signaling culminates in transcriptional regulation is largely unknown. Epigenetic mechanisms are key for integrating external “signals”, e.g., from neighboring cells, into the transcriptome. We have previously reported that ephrinA5 stimulation of immortalized cerebellar granule (CB) cells elicits transcriptional changes of lncRNAs and protein-coding genes. LncRNAs represent important adaptors for epigenetic writers through which they regulate gene expression. Here, we investigate the interaction of lncRNA with protein-coding genes by the combined power of *in silico* modeling of RNA/DNA interactions and respective wet lab approaches, in the context of ephrinA5-dependent regulation of cellular motility. We found that *Snhg15*, a cancer-related lncRNA, forms a triplex structure with the *Ncam1* promoter and interacts with DNMT1. EphrinA5 stimulation leads to reduced *Snhg15* expression, diminished *Snhg15*/DNMT1 interaction and decreased DNMT1 association with the *Ncam1* promoter. These findings can explain the attenuated *Ncam1* promoter methylation and elevated *Ncam1* expression that in turn elicits decreased cell motility of CB cells. Hence, we propose that ephrinA5 influences gene transcription via lncRNA-targeted DNA methylation underlying the regulation of cellular motility.

## Introduction

Cells communicate with the local microenvironment. The perception of those external signals, provided, e.g., by the extracellular matrix (ECM) or cell surface molecules of neighboring cells, is critically involved in regulating cell intrinsic processes that orchestrate cellular proliferation, differentiation, and migration. Apart from proper morphogenesis of tissues and organs, these developmental processes play a key role in tumor initiation and/or progression (Eisenberg et al., 2020; Manzo, 2019; Spill et al., 2016).

The membrane-bound Eph receptors and their cognate ligands, the ephrins, represent signaling molecules that on the one hand orchestrate the development of various tissues including brain structures (Gerstmann et al., 2015; Steinecke et al., 2014; Zimmer et al., 2007; 2008; 2011), and cancer-related aspects on the other hand. The Eph/ephrin system was found to be implicated in numerous types of brain cancer such as glioblastoma and medulloblastoma (Sikkema et al., 2012; Surawska et al., 2004; Uddin et al., 2020). Of note, the expression of ephrinA5 has been found dramatically downregulated in primary gliomas, and the forced expression of *EFNA5* (encoding for ephrinA5) diminishes the tumorigenicity of human glioma cells (Li et al., 2009; Ricci et al., 2020). EPHA2, an Eph receptor known to interact with ephrinA5, has been reported to have not only tumor suppressive but also pro-oncogenic functions (Hamaoka et al., 2016; Wykosky et al., 2005; Wykosky & Debinski, 2008).

Even though the physiological relevance of Eph/ephrin signaling has been well-proven for developmental and cancer-related processes, whether and how the ligand-mediated activation of Eph receptors triggers changes in gene expression that underlie discrete cell physiological responses is greatly unknown. Typically, transcriptional regulation is fine-tuned by epigenetic mechanisms, comprising histone modifications, DNA methylation, and non-coding RNAs (ncRNAs). Apart from functional implications in directing developmental processes, it is widely accepted that dysregulated epigenetic signatures are associated with the initiation and progression of cancer (Anastasiadou et al., 2018; Esteller, 2008; Phillips et al., 2020; Sharma et al., 2010).

DNA methylation, carried out by DNA methyltransferases (DNMTs), is one of the most frequently investigated epigenetic mechanism (Laurent et al., 2010; Liu et al., 2016; Stepper et al., 2017). An important DNA methyltransferase is DNMT1, relevant for *de novo* methylation activity in cancer cells and maintaining the methylation state during proliferation (Al-Kharashi et al., 2018; Gusyatiner & Hegi, 2018). Moreover, DNMT1 was shown to crosstalk with histone modifiers such as histone deacetylases and histone methylases to alter the accessibility of the DNA (Fuks et al., 2000; Symmank et al., 2020). The DNA methylation landscape has been shown to vary dynamically depending on the cell type and developmental stage, and to respond to external signals (Guo et al., 2011; 2014; Skvortsova et al., 2019). DNMT1 function and DNA methylation regulate a broad spectrum of physiological processes, including the migration of neurons (Yildiz & Zimmer-Bensch, 2022) and glioma cells (Hua et al., 2018; Sun et al., 2017). However, whether and how DNMT targets specific gene loci, and induces transcriptionally relevant changes in DNA methylation signatures that elicit physiological responses, is not fully understood. Specifically, to which extent this cascade can be triggered by external signals provided, for instance, by the Eph/ephrin system, remains elusive so far.

We recently provided evidence that the stimulation of cell culture models for medulloblastoma, namely immortalized cerebellar granule (CB) as well as DAOY cells, with ephrinA5, a known tumor suppressor in glioma, has the potential to alter the expression of protein-coding genes and lncRNAs, such as the cancer-relevant lncRNA *SNHG15* (Pensold et al., 2021). In addition to *SNHG15,* abnormal expression of diverse lncRNAs has been implicated in glioma and medulloblastoma molecular pathology (Laneve et al., 2019; Stackhouse et al., 2020). This suggests a functional relevance that needs to be better understood to leverage the potential of lncRNAs as putative therapeutic targets (Ghafouri-Fard et al., 2020; Jiang et al., 2019; Yadav et al., 2021). lncRNAs are known to regulate transcription through interacting with epigenetic writers or erasers (Cabianca et al., 2012; Marchese et al., 2017; Rinn et al., 2007; Zimmer-Bensch, 2019). By forming triplex structures (Kalwa et al., 2016; Kuo et al., 2019; Leisegang et al., 2022; Sentürk Cetin et al., 2019), or during antisense transcription, lncRNAs can promote or prevent the binding of epigenetic modifiers to discrete genomic loci (Zimmer-Bensch, 2019). Here, we aim to test how ephrinA5-dependent signaling regulates gene expression, and cell physiological responses by lncRNA-mediated remodeling of epigenetic signatures.

## Material and Methods

### Cell culture

Cerebellar granule (CB) cells (Fossale et al., 2004) were cultured as previously described (Pensold et al., 2021). Briefly, CB cells were incubated in Dulbecco’s modified Eagle medium (DMEM) with high glucose (#11965084, Gibco), supplemented with 10% fetal bovine serum (FBS) (#S1810, Biowest), 1× GlutaMAX™lJ (#35050038, Thermo Scientific) and 24 mM KCl at 33LJ, 5% CO_2_ and 95% relative humidity. Upon thawing, the medium was additionally supplemented with 100 U/mL penicillin and 100 µg/mL streptomycin until the first passage.

### Treatment with recombinant ephrinA5-Fc

Cells were stimulated with 5 µg/mL of either the recombinant ephrinA5-Fc (#374-EA, Biotechne) or Fc protein (#109-1103, Rockland) as control, both pre-clustered with 10 µg/mL Alexa488-conjugated anti-human IgG (#A11013, Invitrogen) for 30 min at RT.

### Migration assay

Standard TC cell culture plates (#83.3922, Sarstedt) were coated with Geltrex™lJ (#A1413202, Gibco) at a final concentration of 0.2 mg/mL diluted in CB culture medium without phenol red. After a 60 min incubation at 33LJ, the excess medium was removed, and the cells were seeded at a density of 14 cells/mm^2^. After a 24 h incubation, cells were transfected with siRNA oligos at a final concentration of 9 nM by forward lipofection using Lipofectamine 2000© (#11668019, Invitrogen) according to the manufacturer’s protocol. A non-targeting Block-iT Alexa Fluor Red Fluorescent Control siRNA (#14750100, Invitrogen) was utilized as control. 24 h post-transfection, the cells were stimulated with ephrinA5-Fc or Fc protein as control as described in the previous section. After 24 h of ephrinA5-Fc or Fc protein treatment, the cells were imaged for 24 h at 33LJ and 5% CO_2_ using a Leica DMi8 inverted microscope equipped with the Thunder imaging platform. Images were taken every 20 min using a 10× objective and processed with Fiji (ImageJ). Following a minimum intensity z-stack projection, the background noise was reduced using the Basic default plugin, replacing the temporal mean. The corrected image stack was used to create a temporal color code for the first 20 h of imaging to demonstrate the different migration ranges. Next, the background was subtracted using the rolling ball algorithm, and the stack was de-speckled. After converting the stack to 8-bit, the contrast was enhanced, and the stack was binarized using the Yen algorithm. Subsequently, the binarized stack was de-speckled again, and the inconsistencies were fixed using the option “Fill holes”. The wrMTrck plugin was used, as previously described by Sharma et al. (2020), to track cells with a migration time of at least 6 h. The plugin was run with the following parameters: minimum particle size at 180, maximum particle size at 2000, maximum particle velocity at 50, the maximum area change at 400, the minimum track length at 18, and fps at 0.0008. The fastest 50% fraction was used for further analyses.

### Expression analysis via quantitative reverse transcription PCR (RT-qPCR)

Total RNA was purified with the TRIzol™lJ reagent (#15596018, Invitrogen) according to manufacturer’s protocol. Subsequently, samples were treated with RNAse-free DNase I (#EN0521, Thermo Scientific) according to manufacturer’s instructions to eliminate possible genomic DNA contaminants. cDNA synthesis was performed by reverse transcription using the iScript cDNA Synthesis Kit (#1708890, Bio-Rad). Quantitative real-time PCR (qPCR)

reactions were performed with 10 ng cDNA of each sample and the PowerUP SYBR Green qRT-PCR Kit (#A25741, Applied Biosystems) using the CFX96 thermocycler (Bio-Rad). Primer sequences are listed in Supplementary Table S1. Data analysis was performed via the previously described ΔΔCt method (Livak & Schmittgen, 2001) using the reference gene *Atp5bp*. Normalized expression levels were calculated relative to control-Fc-treated samples.

### Chromatin immunoprecipitation (ChIP)

24 h following the ephrinA5-Fc or Fc stimulation, 1.5×10^6^ cells were lysed with digestion buffer (50 mM Tris-HCl pH 8.0, 1 mM CaCl_2_, 0.2% (v/v) Triton X-100, 1% protease inhibitor cocktail (#I3786, Merck)). Chromatin was enzymatically sheared for 5 min at 37°C with 0.2 mU/μL Micrococcal nuclease (MNase) (#N3755, Merck), and stopped with MNase stop buffer (110 mM Tris-HCl pH 8.0, 55 mM EDTA). After adding 2x RIPA buffer (280 mM NaCl, 1.8% (v/v) Triton X-100, 0.2% (v/v) SDS, 0.2% (v/v) sodium deoxycholate, 5 mM EGTA, 1% protease inhibitor cocktail), the samples were centrifuged for 15 min at 4°C and 21,130×g. Following the centrifugation, 1% (v/v) of the supernatant was used as input control whereas 20% (v/v) was used for each immunoprecipitation (IP). The input control was incubated in TE buffer (10 mM Tris-HCl pH 8.0, 1 mM EDTA) supplemented with 40 mU/µL proteinase K (#P4850, Merck) for 2 h at 55°C and 1200 rpm. Per IP, 25 μL of protein A-coupled Dynabeads (#10001D, Invitrogen) were prepared by washing them twice and resuspending to the original volume with 1x RIPA buffer (10 mM Tris-HCl pH 8.0, 1 mM EDTA, 140 mM NaCl, 1% (v/v) Triton X-100, 0.1% (v/v) sodium deoxycholate, 0.1% (v/v) SDS, 0.1% protease inhibitor cocktail). Then, the IP samples were pre-clearead with 10 μL of Dynabeads for 1 h at 4°C with rotation. Following the pre-clearing, the beads were discarded, and the IP samples were incubated overnight with 40 mg/mL rabbit anti-DNMT1 (#70201, BioAcademia), mouse anti-H3K27me3 (#Ab6002, Abcam) or normal rabbit IgG (#12-370, Merck) antibody at 4°C with rotation. After the overnight incubation, 10 μL of Dynabeads were added to the IP samples, followed by a 3 h incubation with rotation at 4°C. Subsequently, the antibody-bound beads were washed five times with 1x RIPA buffer, once with LiCl wash buffer (250 mM LiCl, 10 mM Tris-HCl pH 8.0, 1 mM EDTA, 0.5% (v/v) Igepal CA-630, 0.5% (v/v) sodium deoxycholate, 0.1% protease inhibitor cocktail), and once with TE buffer. Following the final wash, the beads were resuspended in TE buffer with 40 mU/µL proteinase K (#P4850, Merck) and incubated for 2 h at 55 °C and 1200 rpm. The DNA from the input control and IP samples was isolated using the ChIP DNA Clean and Concentrator Kit (#D5205, Zymo Research) according to manufacturer’s guidelines. ChIP-qPCRs were performed using the isolated ChIP-DNA and input control DNA as templates and the PowerUP SYBR Green qRT-PCR Kit (#A25741, Applied Biosystems) on the CFX96 thermocycler (Bio-Rad). The primer sequences are listed in Supplementary Table S1. Data analysis was performed by a double normalization, first against the input control to calculate recovery and then against IgG to calculate the fold enrichment.

### UV-crosslinked immunoprecipitation (CLIP)

After 24 h of ephrinA5-Fc and Fc control treatment, cells were washed once with pre-warmed 1x Dulbecco’s phosphate-buffered saline (DPBS) (#14190-094, Gibco). Next, 6 mL ice-cold DEPC-treated PBS was applied to the cells which were subsequently irradiated with 150 mJ/cm^2^ at 254 nm for 40 s, harvested with a cell scraper, gently homogenized, and transferred into microtubes. The cells were pelletized at 4LJ and 22,000×g for 30 s and lysed with the pre-cooled lysis buffer (50 mM Tris-HCl (pH 7.4), 100 mM NaCl, 1% (v/v) Tergitol, 0.1% (v/v) SDS, 0.5% (v/v) sodium deoxycholate) supplemented with 1% protease inhibitor cocktail and 2% RiboLock RNase Inhibitor (#EO0381, Thermo Scientific). The RNA was sheared, and the DNA was degraded for 3 min at 37LJ and 300 rpm using 45 mU/mL MNase (#N3755, Merck) and 36 U/mL DNase (#EN0521, Thermo Scientific). The reaction was stopped using the MNase stop buffer (110 mM Tris-HCl pH 8.0, 55 mM EDTA). The samples were centrifuged for 10 min at 4 LJ and 22,000×g, and the supernatant was transferred into RNase-free microtubes. 1% (v/v) of the sample was isolated as input control, mixed with 500 µL TRIzol (#15596018, Invitrogen), snap frozen on dry ice and stored at -20LJ until further use. Per immunoprecipitation, 25 mg/mL of rabbit anti-DNMT1 (#70201, BioAcademia), rabbit anti-EZH2 (#5246S, Cell Signaling) or normal rabbit IgG (#12-370, Merck) antibody were pre-incubated with washed Dynabeads (#10001D, Invitrogen) for 1h at RT to pre-coat the beads with antibodies. The IgG pulldown was applied to differentiate signal from noise due to unspecific binding of lncRNAs to rabbit epitopes (Lee & Ule, 2018). 33% (v/v) of the sheared RNA samples were added to the different antibody-bead mixtures and rotated for 2h at 4LJ to allow for the antibodies to bind to their target protein. Next, the samples were washed twice with high-salt buffer (50 mM Tris-HCl (pH 7.4), 1 M NaCl, 1 mM EDTA, 1% (v/v) Tergitol, 0.1% (v/v) SDS, 0.5% (v/v) sodium deoxycholate) followed by two washes with wash buffer (20 mM Tris-HCl (pH 7.4), 10 mM MgCl_2_, 0.2% (v/v) Tween-20) at 4LJ, with each washing step lasting for 1 min with rotation. After discarding the supernatant, the beads were resuspended in wash buffer supplemented with 20 mU/µL proteinase K (#P4850, Merck) and incubated with shaking for 20 min at 37□ and 300 rpm. Afterwards, the CLIP-RNA was purified alongside the input control with TRIzol™lJ reagent (#15596018, Invitrogen) according to manufacturer’s protocol and used as template for the quantitative reverse transcription PCR. The CLIP-RT-qPCRs were performed using the SuperScript™lJ III Platinum™lJ SYBR™lJ Green One-Step kit (#11736-051, Invitrogen) on the CFX96 thermocycler (Bio-Rad). The primer sequences are listed in the Supplementary Table S1. RNA recovery was calculated via normalization to the total amount of RNA per experiment and condition.

### DNA Methylation Profiling

24 h after ephrinA5-Fc or Fc stimulation, CB cells were harvested and their DNA was extracted using the PureLink® Genomic DNA Mini Kit (#K1820, Invitrogen). The samples were then treated with proteinase K and RNase A supplied in the kit. The DNA methylation profiling was carried out using the Infinium Mouse Methylation BeadChip (Illumina) according to the manufacturer’s standard protocol. 500 ng of genomic DNA was bisulfite converted using the Zymo EZ-96 DNA Methylation kit (Zymo Research, Irine, CA, USA). Subsequently, the bisulfite converted DNA samples were amplified, fragmented, purified, and hybridized onto the BeadChip array following the manufacturer’s protocol. The arrays were washed and scanned using the Illumina iScan System. Mouse Methylation BeadChips were processed at Life & Brain (L&B) Genomics, Bonn.

### Differential DNA Methylation Analysis

Differentially methylated regions (DMRs) were detected using the R packages Enmix (Xu et al., 2016), sva (Leek, 2007) and minfi (Aryee et al., 2014). The raw idat files were loaded by Enmix::readidat() together with Illumina’s Infinium mouse-methylation manifest file (v.1.0). For background correction, dye bias correction, inter-array normalization and probe type bias correction we applied Enmix::mpreprocess() on the raw idat data setting the parameters qc and impute to TRUE. It returns a matrix of preprocessed methylation beta values. As a second preprocessing step we used sva::ComBat() to mitigate the batch effect contained in these beta values which was introduced by different experiment runs under inevitably different conditions. The experiment run ID was set as the batch variable. Eventually, DMRs were identified by calling minfi::dmpFinder() on the preprocessed beta values. The parameter pheno was set to the respective cell conditions (for each sample either ctrl-Fc or efnA5-Fc) and the parameter type to =”categorical”. CpGs for which the dmpFinder result indicates a p-value of 0.05 or smaller were considered significant DMRs. If not indicated otherwise, default parameters have been passed to the applied R functions.

### In silico simulation of RNA/DNA interactions

For *Adamts14,* the sequence-based predictions (method described in Pensold et al., 2021; sequences presented in Supplementary Table S2) suggested two slightly different alternatives for the 15-nucleotide binding mode, which were used to generate the two models, *Adamts14*-1 and *Adamts14*-2. For *Ncam1,* a 15-nucleotide sequence (*Ncam1*) as well as an extended sequence (*Ncam1*-ext) with additional base pairs (see Supplementary Table S2) were chosen. The latter was introduced, as our preliminary evaluation revealed the binding site boundaries to be too narrow in this case and to cause artefactual strand separation. The extended version allowed us to evaluate the impact of termini fluctuations on the stability of the system in the simulations. To maintain comparability, we only evaluated sections that are also present in the non-extended *Ncam1* model.

All molecular dynamics simulations were carried out with the GROMACS simulation package (version 2021.4) using the AMBER-parmBSC1 force field (Ivani et al., 2016) and TIP4P-D water model (Piana et al., 2015) in a rhombic dodecahedral box with periodic boundaries under standard conditions (300 K, 1 bar). We parameterized a protonated cytosine and used it for all cytosines that were not terminal. Potassium chloride, sodium chloride, and magnesium ions were added to the system. For the ions, the Joung and Cheatham parameters (Joung & Cheatham III, 2008) were used. The concentrations are tuned to mimic the cellular environment with a sodium chloride concentration of 0.01 M and potassium chloride concentration of 0.1 M while magnesium ions were introduced to neutralize the total charge of the system. All ions were placed randomly in the simulation box.

To prepare for the simulation, each system underwent the following procedure: The potential energy of the system was minimized to eliminate clashes and bad contacts by using steepest descent energy minimization followed by conjugate gradient as implemented in GROMACS (Abraham et al., 2015). The initial minimization was followed by three preparatory steps: First the system was heated up to 300 K by gradually increasing the temperature of the system from 0 to 300 K in 10 steps, lasting 1 ns each, using a Berendsen thermostat. Next, a simulation using an NVT (constant number, volume, and temperature) ensemble was conducted for 10 ns using position restraints with a force constant of 1000 kJ/(mol*nm^2^) applied to all heavy atoms. Finally, a simulation in an NPT (constant number, pressure, and temperature) ensemble was conducted at 1 bar and 300 K for 10 ns. Velocity rescaling was used for temperature coupling with a time constant of 0.1 ps in order to ensure correct temperature fluctuations. For simulations at constant pressure, we used the Parrinello-Rahman pressure coupling algorithm (Parrinello & Rahman, 1981) with a time constant of 2 ps. Afterwards at least 600 ns were simulated under NPT ensemble conditions with an integration step of 2 fs. All bonds were constrained using the LINCS algorithm (Hess, 2008). A cutoff of 10 Å was used for Lennard-Jones and short-range Coulombic interactions and the particle mesh Ewald (PME) method was used for long-range electrostatic interactions with a grid spacing of 0.16 nm and an interpolation order of 4. The short-range Lennard-Jones interactions were handled by using the grid system for neighbor searching. The cut-off distance for the neighbor list was 1.2 nm.

For visualization of the trajectory, we used VMD 1.9.3. For the analysis, we used VMD, tools provided by GROMACS, and our own scripts. Before analyzing the geometric properties of the trajectory, we eliminated periodic jumps and centered the solute using gmx trajconv.

## Results

### *Snhg15* interaction with DNMT1 is diminished upon ephrinA5-Fc stimulation

We demonstrated earlier that ephrinA5-Fc stimulation of CB cells modulates the expression of protein-coding genes as well as of non-coding RNAs, including the cancer-associated lncRNA *Snhg15* (Pensold et al., 2021). Its human ortholog *SNHG15* has cancer- and metastasis-promoting functions, linked to poor survival in numerous human malignancies (Tong et al., 2019). In addition to its well-documented function as a competitively endogenous RNA (ceRNA) consisting of sponging miRNAs in human cancers (Wu et al., 2018), *SNHG15* was reported to act in the nucleus in concert with EZH2 (Enhancer of zeste homolog 2), which is known to catalyze repressive trimethylation at histone 3 (H3K27me3) (Z. Ma et al., 2017). We found the expression of *Snhg15* to be reduced upon ephrinA5-Fc stimulation in CB and DAOY cells (Pensold et al., 2021). Using computational approaches, we predicted *Snhg15* to be capable of interacting with the promoters of 19 protein-coding genes that were increased in expression upon ephrinA5-Fc stimulation (Pensold et al., 2021). These interactions were presumed to be driven by a predicted DNA-binding domain (DBD) in *Snhg15*, localized in nucleotide (nt) positions 1896-1925 (Pensold et al., 2021). Here, we asked whether these ephrinA5-triggered transcriptional changes of protein-coding genes are facilitated through *Snhg15*-mediated actions. These could be alterations of repressive epigenetic marks, since lncRNAs were reported to recruit or to interact with epigenetic writers including DNMTs (Chalei et al., 2014; Guil et al., 2012; Wang et al., 2015).

First, we verified whether *Snhg15* indeed interacts with epigenetic writers of repressive chromatin states, and whether this interaction is changed upon ephrinA5-Fc stimulation. To this end, we performed UV-mediated crosslinking of RNA with proteins in CB cells, followed by the immunoprecipitation of the protein of interest (CLIP). We chose to profile putative interactions of *Snhg15* with two major repressive epigenetic writers: EZH2, as the main enzyme of the PRC2 (Polycomb repressive complex 2) that catalyzes repressive trimethylations at H3K27 residues (Cao et al., 2008; Kuzmichev et al., 2002; Margueron & Reinberg, 2011) and was described to act in concert with *Snhg15* (Z. Ma et al., 2017); as well as DNMT1, one of the main DNA methyltransferases which catalyzes DNA methylation and acts mainly repressive on gene regulation (Laurent et al., 2010; Liu et al., 2016; Stepper et al., 2017; Zemach et al., 2010). Both proteins have been frequently implicated in transcriptional dysregulation which is a hallmark of glioma and medulloblastoma pathogenesis (Chen et al., 2021; Hervouet et al., 2010; Miele et al., 2017; Pócza et al., 2016; Rajendran et al., 2011; Stazi et al., 2019; Zhang et al., 2020). To investigate the potential interaction of EZH2 and/or DNMT1 with *Snhg15*, we performed antibody-mediated pulldown of EZH2- and DNMT1-bound RNA using CLIP. Three different primer pairs covering distinct positions and isoforms of *Snhg15* (Figure 1a, Supplementary Figure S1) were used for RT-qPCR-based analysis of the co-immunoprecipitated RNA. From the four known murine isoforms, only two isoforms (isoform 202 and 203) display the predicted DNA binding domain (Figure 1a). Both were covered by primer pair *SP1*. Primer pair *SP2* is specific for isoform 202, while primer pair *SP3* only detects isoform 203 (Figure 1a).

**Figure 1:**
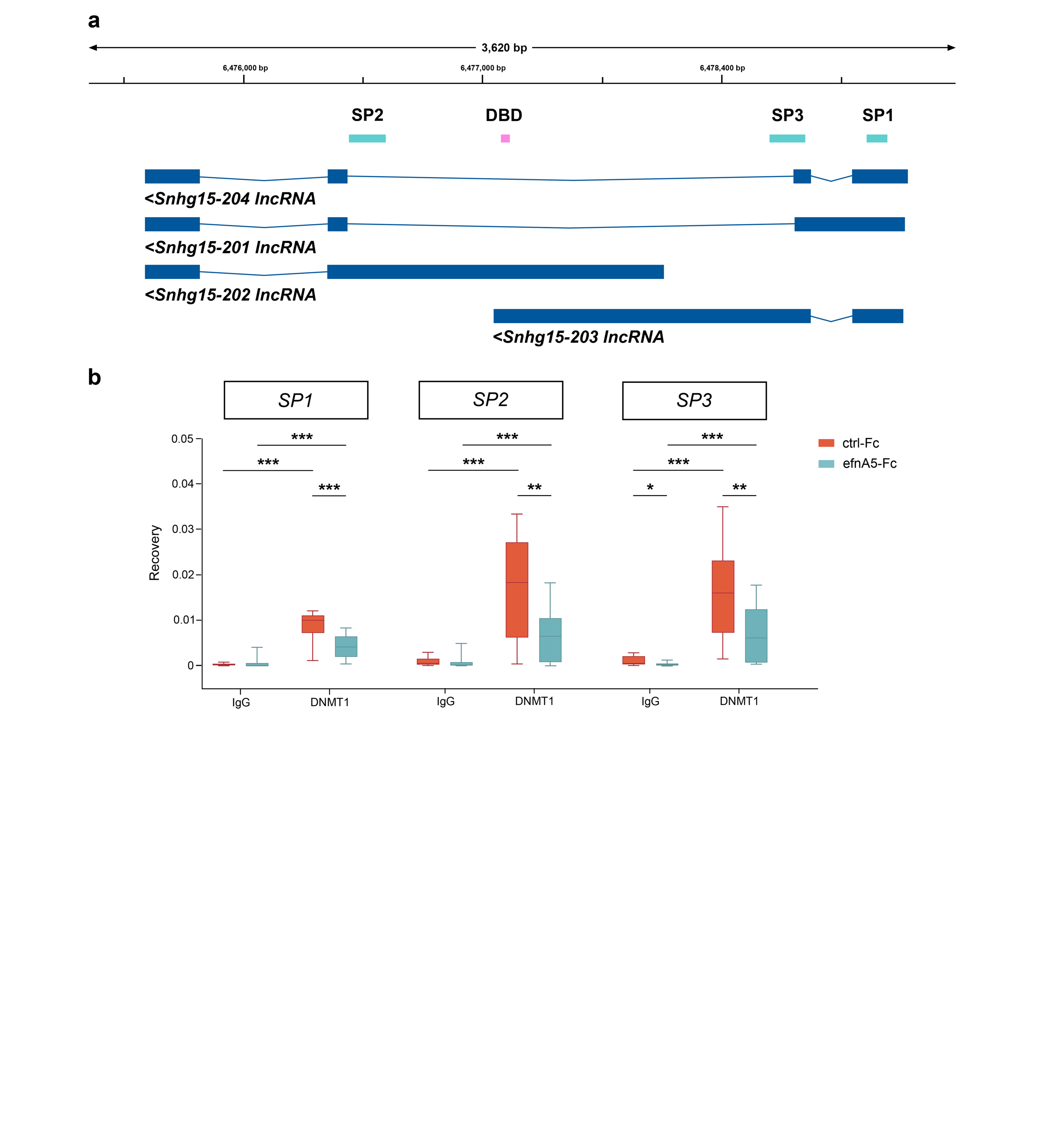
*Snhg15* is associated with DNMT1 in an ephrinA5-dependent manner in CB cells. (a) Location of the three *Snhg15* amplicons used to amplify CLIP-RNA (from left to right: *SP2*, SP3, *SP1*) and the DNA-binding domain (DBD) location. (b) RNA recovery for IgG and anti-DNMT1 antibody CLIP samples in CB cells (N = 5 biological replicates, min-to-max box plots). For amplicon positions *SP1*, *SP2*, and *SP3*, CLIP clearly indicated a (functional) association of DNMT1 with *Snhg15,* and a reduction of the amount of bound *Snhg15* RNA after ephrinA5-Fc stimulation. Significances were determined with two-tailed Student’s *t*-test. Significance levels: *p* value < 0.05 *; *p* value < 0.01 **; *p* value < 0.001 ***. ctrl-Fc: control-Fc, efnA5-Fc: ephrinA5-Fc, CLIP: UV cross-linking and immunoprecipitation.

CLIP experiments using the EZH2 antibody did not yield an RNA recovery above background noise, neither for control conditions (control-Fc-treated cells) nor for the samples treated with ephrinA5-Fc (Supplementary Figure S1). In contrast, pulldown using a DNMT1-specific antibody resulted in a significant RNA recovery compared to IgG-pulldown experiments at all tested loci (Figure 1a and b), indicative of DNMT1 binding to *Snhg15*. Interestingly, ephrinA5-Fc treatment significantly reduced the relative amount of DNMT1 association to *Snhg15* fragments at the tested loci (Figure 1b). Together, these data propose that *Snhg15* directly interacts with DNMT1, and that this interaction is diminished by ephrinA5 stimulation.

### EphrinA5-Fc stimulation impacts DNA methylation at promoters of *Snhg15* target genes such as *Ncam1*

DNMT1 catalyzes DNA methylation, which is often associated with gene repression (Laurent et al., 2010; Li et al., 1992; Liu et al., 2016; Zemach et al., 2010). Thus, we next aimed to analyze, whether ephrinA5-Fc stimulation of CB cells induces changes in DNA methylation. To this end, DNA samples from CB cells treated with ephrinA5-Fc and Fc-control were analyzed for changes in CpG methylation using the Infinium Mouse Methylation BeadChip array. We detected numerous genes with altered CpG levels, indicating that ephrinA5-Fc stimulation triggers changes in DNA methylation (Figure 2a and b, Supplementary Table S3), which could explain the transcriptional changes observed earlier after ephrinA5-Fc treatment (Pensold et al., 2021).

**Figure 2:**
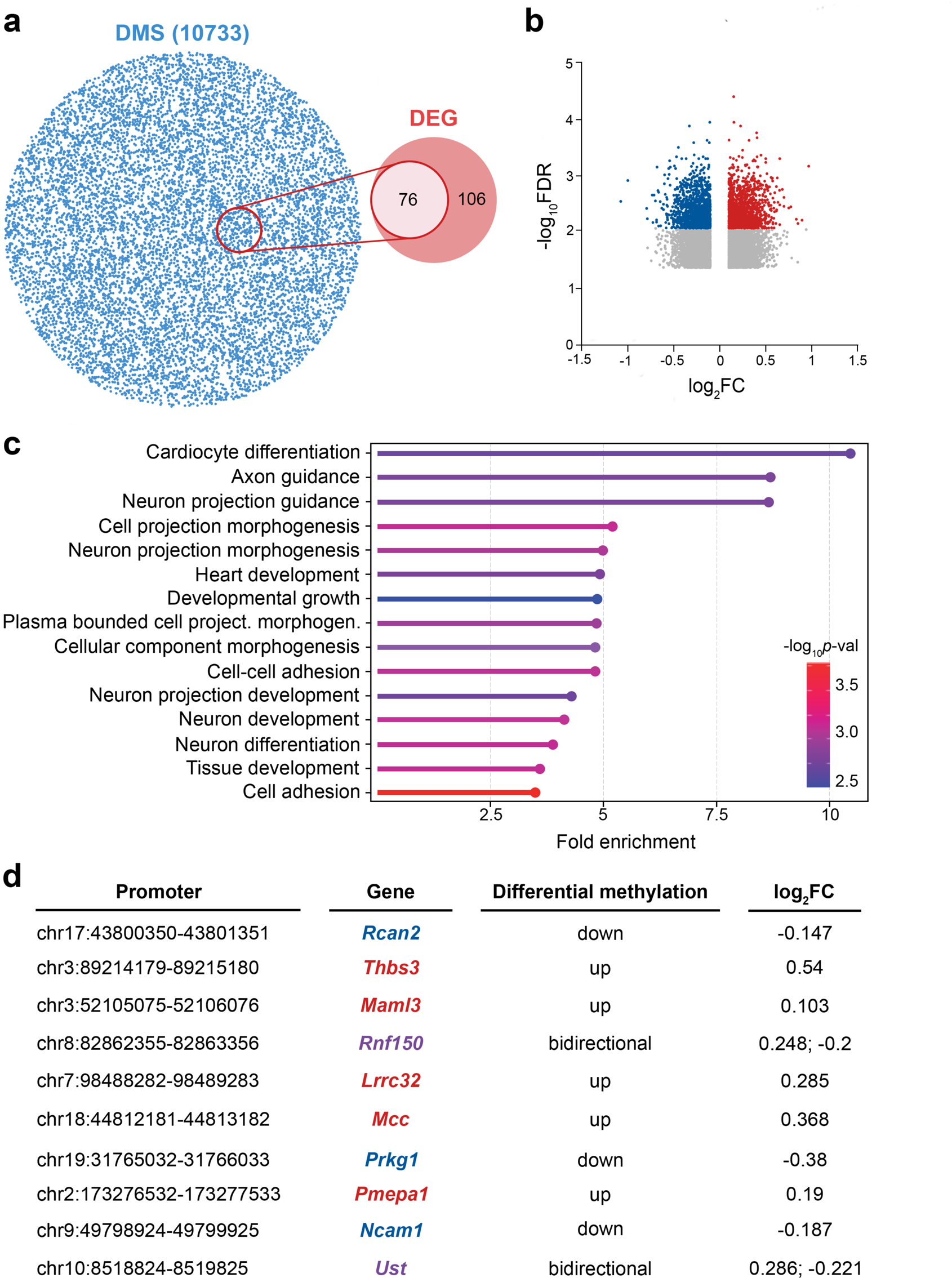
EphrinA5-Fc stimulation induces changes in DNA methylation signatures, including cell adhesion-related genes. (a) Venn diagram shows overlap between differentially expressed genes (DEG) and differentially methylated sites (DMS) in ephrinA5-Fc-treated versus control-Fc-treated CB cells. (b) Volcano plot displays loci with significantly increased and decreased levels of DNA methylation, shown in red or blue, respectively, while differences below -log_10_*p*-val after false discovery rate (FDR) correction are depicted in gray. Loci changed below 0.1 log_2_FC (fold change) are excluded from the plot. (c) Gene Ontology analysis (GO) reveals enrichment for several GO terms related to cell adhesion and neuronal development; fold enrichment represented by the x-axis and -log_10_*p*-val is encoded by the color gradient. (d) Table lists all protein-coding genes with predicted binding sites for *Snhg15* that are upregulated (RNA-seq; Pensold et al., 2021) and simultaneously display DNA methylation changes 24 h after ephrinA5-Fc treatment. Genes with reduced methylation are presented in blue font, red font indicates increased methylation after ephrinA5-Fc treatment, while for genes with purple font CpG sites with increased as well as decreased methylation levels were detected.

In support of these findings, about 42% of the differentially expressed genes displayed concomitant changes in DNA methylation (Figure 2a). Among these genes that were both changed in transcription and in CpG methylation after ephrinA5-Fc stimulation, we determined a significant enrichment of cell adhesion-related genes by Gene Ontology (GO) analysis (Figure 2c).

When focusing on the 19 genes upregulated after ephrinA5-Fc treatment and predicted to be bound by *Snhg15* via triple helix-mediated RNA-DNA interaction (Supplementary Table S4; Pensold et al., 2021), 10 of these genes showed significant alterations in the methylation of CpG sites. Methylation changes were located at intragenic loci (introns and exons), transcription start sites (TSSs) or upstream of the TSSs, with decreased as well as increased methylation levels in response to ephrinA5-Fc treatment (Figure 2d, Supplementary Table S3). Methylation at the TSSs is usually associated with gene repression (Jones, 2012). According to our hypothesis, *Snhg15* interacts with DNMT1 as a repressive epigenetic writer, thereby recruiting DNMT1 to discrete gene loci, e.g., transcription start sites, and in concert leads to gene repression. Since the interaction of *Snhg15* and DNMT1 was diminished by ephrinA5-Fc stimulation, we were screening for genes with reduced CpG methylation close to their TSS among the 10 remaining genes (Figure 2d). For *Rcan2*, *Prkg1*, and *Ncam1*, we detected an ephrinA5-Fc treatment-induced decrease in CpG methylation levels close to TSSs. Of note, *Ncam1* encodes the well-known adhesion protein NCAM1 (neural cell adhesion molecule 1), a crucial key player in not only neuronal but also cancer cell migration (Prag et al., 2002). For various cancer types, elevated expression of *Ncam1*/*NCAM1*, like we observed after ephrinA5-Fc stimulation, is associated with reduced tumor cell migration and better prognosis (Edvardsen et al., 1994; Guan et al., 2020). In line with this, we detected in patient data that increased expression levels of *NCAM1* in low-grade glioma is associated with improved survival rates (Supplementary Figure S2a).

### Increased *Ncam1* expression level contributes to the ephrinA5-Fc triggered decrease in CB cell motility

In agreement with the inverse correlation of *NCAM1* expression with tumor cell migration (Edvardsen et al., 1994; Guan et al., 2020), we found reduced motility of CB cells, when analyzing their migration *in vitro* 24 h after ephrinA5-Fc stimulation (Figure 3a and b). This time point was chosen since the elevated expression and diminished promoter methylation of *Ncam1* were detected here (Figure 2d). To verify, whether the ephrinA5-Fc-induced increase in *Ncam1* transcript levels accounts for the motility changes of CB cells, we performed live cell imaging experiments of cells 24 h after ephrinA5-Fc or control-Fc stimulation, with a preceding siRNA-mediated knockdown of *Ncam1* (see Supplementary Figure S2b for validation of knockdown efficiency). Indeed, the ephrinA5-Fc stimulation-triggered reduction of the migratory speed of CB cells compared to control-Fc conditions was rescued by *Ncam1* siRNA application (Figure 3a and b). Of note, ephrinA5 is known to bind EphA2 which we’ve previously demonstrated to be highly expressed in CB cells (Pensold et al., 2021). To test, whether the ephrinA5-Fc triggered effect on cell motility in CB cells was dependent on EphA2, we knocked down the *EphA2* expression with target specific siRNA oligos prior to ephrinA5-Fc or control-Fc stimulation of cells (knockdown efficiency for the EphA2 is depicted in Supplementary Figure S3c). As a matter of fact, we found the motility reducing effect of ephrinA5-Fc stimulation to be reversed to control levels after siRNA-mediated knockdown of *EphA2* (Supplementary Figure S3a and b). Together, these data propose that the ephrinA5-Fc triggered raise in *Ncam1* expression accounts for the impaired motility of CB cells, and that this mechanism is mediated by the EphA2 receptor.

**Figure 3:**
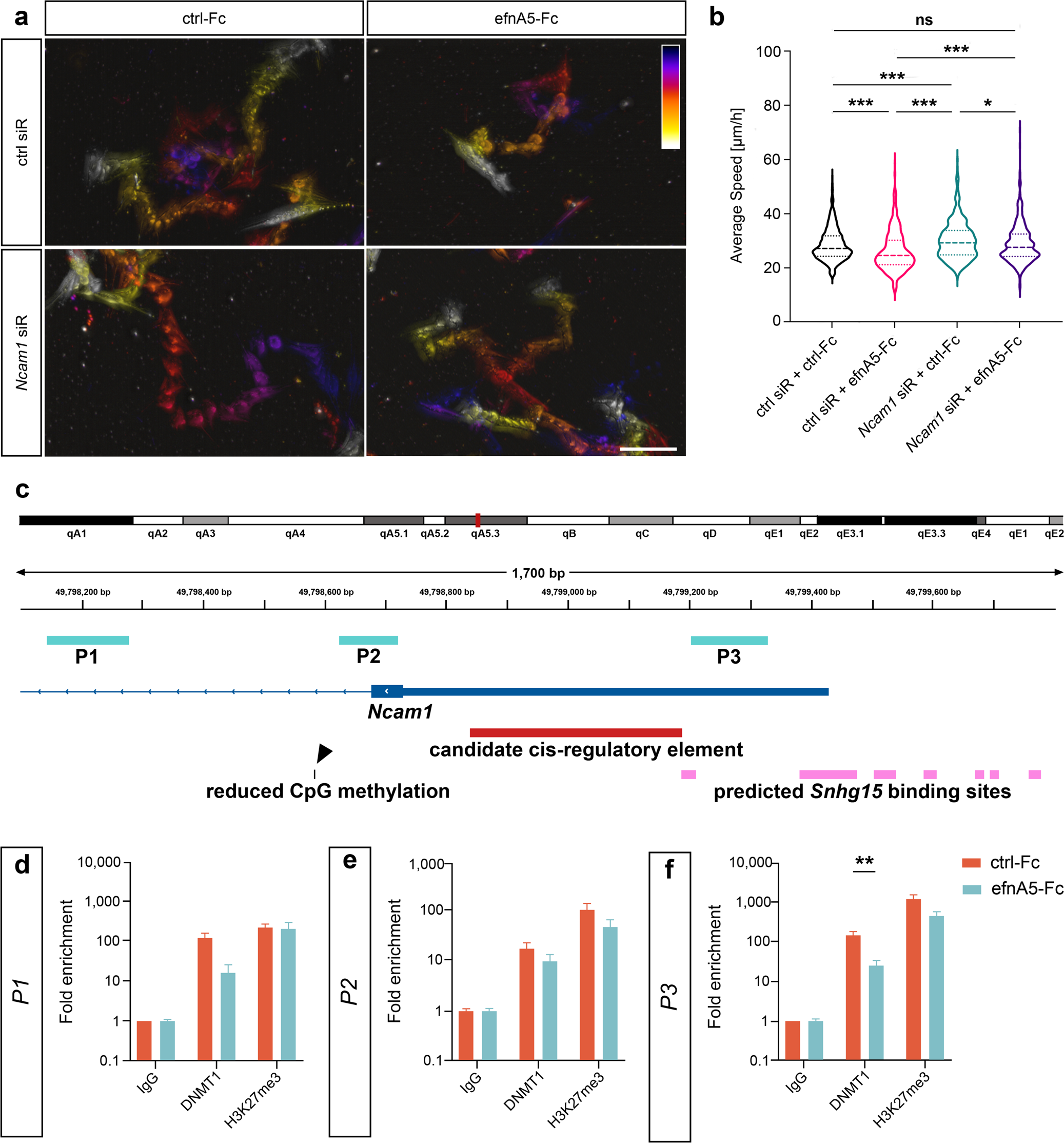
EphrinA5 stimulation leads to reduced association of DNMT1 to *Ncam1* and diminished motility of CB cells. (a-b) The motility of CB cells is significantly reduced upon stimulation with ephrinA5-Fc, which can be rescued by a preceding knockdown of *Ncam1*. (a) Temporal color-coded migratory distance over 20 h of imaging. The starting point of migration for each cell is shown in dark blue and the end point in white. (b) Quantitative analysis of average migratory speed (n = 557 for ctrl siR + ctrl-Fc, n = 455 for ctrl siR + efnA5-Fc, n = 481 for *Ncam1* siR + ctrl-Fc, n = 495 for *Ncam1* siR + efnA5-Fc, N = 4 biological replicates). (c-f) Native ChIP revealed decreased enrichment of DNMT1 in the *Ncam1* promoter region close to putative *Snhg15* binding sites. (c) Genomic map depicting the promoter region of the murine *Ncam1* gene. Regions targeted by the primer pairs *P1*, *P2* and *P3* are shown in turquoise, the promoter in dark blue, candidate *cis*-regulatory elements in red, CpG site with a significantly reduced methylation level in black, and putative *Snhg15* binding sites in pink. (d-f) ChIP-qPCR analysis using anti-DNMT1 and anti-H3K27me3 antibodies normalized against the input material and IgG (N=4 biological replicates). Significances were determined with one-way ANOVA (b) and two-tailed Student’s *t*-test (d-f). Significance levels: *p* value < 0.05 *; *p* value < 0.01 **; *p* value < 0.001 ***. Scale bar: 100 μm. ctrl: control. efnA5: ephrinA5. siR: siRNA.

### *Ncam1* is a potential target for *Snhg15*-mediated recruitment of DNMT1 and DNMT1-dependent DNA methylation

As we found decreased DNA methylation levels close to the TSS of *Ncam1* after ephrinA5-Fc stimulation of CB cells (Figure 2d; 3c), we next aimed to elucidate whether DNMT1 binds to the *Ncam1* locus in an ephrinA5-Fc stimulation-dependent manner. Of note, in this current study we have shown that ephrinA5-Fc treatment diminishes the interactions of DNMT1 with *Snhg15* (Figure 1a and b). Hence, we next performed native chromatin immunoprecipitation (ChIP) in CB cells treated with ephrinA5-Fc and control-Fc for 24 h prior to sample collection. A target-specific antibody was applied to pull down DNMT1, and qPCR was performed to quantitatively assess the co-immunoprecipitated DNA fragments. To detect enrichment on *Ncam1*, we used primer pairs targeting regions downstream (*P1* and *P2*) and upstream of the promoter (*P3*). For regions covered by *P1* and *P2* we did not detect any significant changes in the enrichment with DNMT1 upon stimulation with ephrinA5-Fc (Figure 3c-e). The region covered by primer pair *P3*, located in close proximity to putative *Snhg15* binding sites and next to a candidate *cis*-regulatory element, presented a diminished association with DNMT1 after ephrinA5-Fc stimulation (Figure 3f). In contrast to the results for *Ncam1*, for *Adamts14*, no significant changes in DNMT1 enrichment were detected for a locus next to the candidate cis-regulatory element (upstream of its promoter) and in proximity to a putative *Snhg15* binding site after ephrinA5-Fc stimulation (Supplementary Figure S4). In line with this, no methylation changes at CpG sites were observed for *Adamts14* (Supplementary Table S3). This data points to a DNA methylation-independent upregulation of *Adamts14* after ephrinA5-Fc stimulation. No alterations in H3K27me3 occupation were observed for any of the targeted regions (Figure 3d-f; Supplementary Fig S4).

In sum, the *Ncam1* promoter region potentially serves as a specific target for *Snhg15*-mediated recruitment of DNMT1 and DNMT1-dependent DNA methylation, which is diminished upon ephrinA5-Fc stimulation. These results are in line with increased expression and reduced methylation levels of *Ncam1*.

### *In silico* modelling confirms potential of *Snhg15* triple helix interaction at *Ncam1* promoter

After having shown, that *Snhg15* binds DNMT1, and DNMT1 associates with the *Ncam1* promoter, with a reduction of both interactions after ephrinA5-Fc treatment, we next aimed to assess whether *Snhg15* forms a sequence-specific triple helix with the *Ncam1* locus. To this end, we performed *in silico* modeling in atomistic detail using molecular dynamics simulations on the sequences of *Snhg15* DBD and the previously predicted *Snhg15* binding sites at the *Ncam1* promoter sequence (Supplementary Table S2). In addition, the predicted triple helix formed by *Snhg15* binding was modeled for the *Adamts14* promoter sequence. In contrast to *Ncam1*, the *Adamts14* locus did not show altered methylation levels and DNMT1 association in the promoter region (Supplementary Table S3).

The sequences chosen for atomistic modelling were based on predicted triple helices for *Ncam1* and *Adamts14* (Kuo et al., 2019; Pensold et al., 2021) (Supplementary Table S2). Specifically, we have two slightly different sequences for *Adamts-14* (*Adamts14-1* and *Adamts14-2*, hereafter, see Methods for details) and for *Ncam1* (*Ncam1* and *Ncam1-ext*, hereafter, see Methods for details). All of the modeled systems are parallel triple helices, where the RNA strand (pyrimidine strand) is oriented in parallel with the purine DNA strand (Buske et al., 2011). For the analysis, we focused on the converged parts of the trajectories (i.e., the last 400 ns, see Supplementary Figure S5).

We first evaluated the interaction energy between the RNA and the DNA as a sum of Lennard-Jones (LJ) and Coulomb (CB) energies. We observed that the RNA strand interacts more extensively with DNA when using the *Adamts14*-1 sequence compared to *Adamts14*-2 (Figure 4b). The same is true for *Ncam1*-ext with respect to *Ncam1* (Figure 4b).

**Figure 4:**
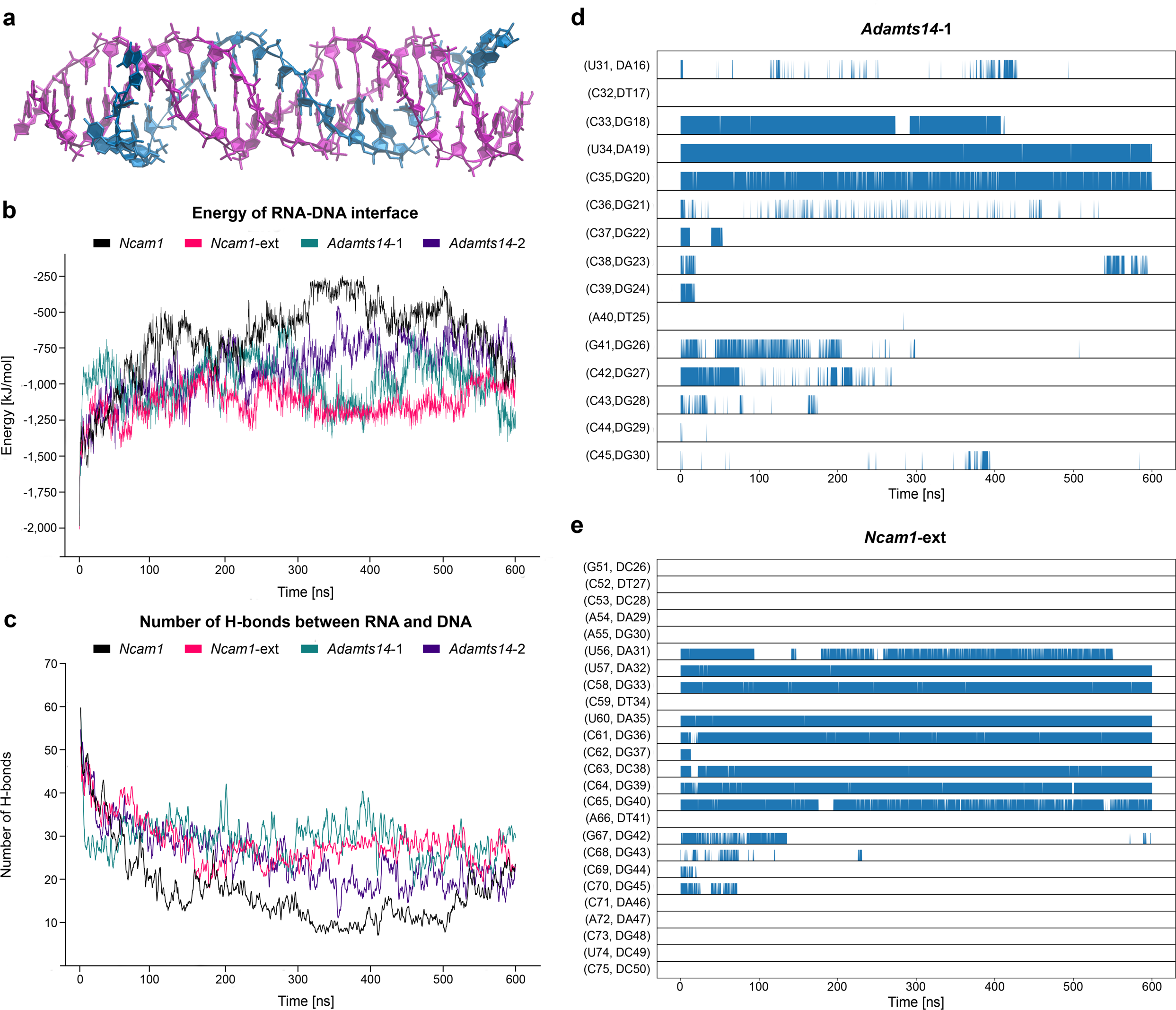
*In silico* modeling of *Snhg15* binding to *Ncam1* and *Adamts14* promoter regions. (a) Snapshot of *Snhg15* triple helix formation with the extended *Ncam1* sequence (*Ncam1*-*ext*) during MD simulations. (b) Total interaction energy (LJ + CB) between DNA and RNA with phosphate and sugar backbone included as a function of time. When comparing *Ncam1* and *Ncam1*-*ext*, we only considered the sequence without the extension (clipped *Ncam1*-*ext* sequence, see Supplementary Table S2) to maintain comparable energies for both models. (c) Number of hydrogen bonds between RNA and DNA. Please note that the hydrogen bonds counts have been smoothed using a running average with a window size of 5, as the curves would otherwise overlap too much. Again, we only considered the sequence without the extension (clipped *Ncam1*-*ext* sequence) to compare *Ncam1* and *Ncam1*-*ext*. (d) Occurrence of *in-register* hydrogen bonded pairs between the RNA and the DNA purine strand in *Adamts14*-*1* along the simulation trajectory. Labels starting with a “D” indicate the DNA residue of the pair. (e) Occurrence of in-register hydrogen bonded pairs between the RNA and the DNA purine strand in *Ncam1*-*ext* (clipped) along the simulation trajectory. Note that predicted interactions get more stable over time as indicated by less noise in the lower lanes. Labels starting with a “D” indicate the DNA residue of the pair. LJ: Lennard-Jones. CB: Coulomb.

We therefore tried to rationalize these trends by breaking down the molecular interactions between the nucleic acid strands. These can be classified in mainly three categories: stacking interactions between the stacked bases in each strand, cross-term interactions between non-adjacent bases across different strands, and hydrogen bonds between the strands. By evaluating the stacking and the cross-term energies of the four systems, we observed that these are comparable between *Adamts14*-*1* and *Adamts14*-*2* (Supplementary Figure S6), as well as between *Ncam1* and *Ncam1*-*ext* (Supplementary Figure S7). This suggests that the difference in stability between the systems are mainly due to the hydrogen bonding between the strands. We indeed found that *Adamts14*-1 and *Ncam1*-*ext* are both able to maintain on average around 27 hydrogen bonds between the RNA and the DNA helix (Figure 4c), while *Adamts14*-*2* and *Ncam1* showed only around 20 and 15 hydrogen bonds, respectively. From this data, we conclude that *Adamts14*-*1* and *Ncam1*-*ext* are the better models for simulating triple helix interaction dynamics compared to *Adamts14*-*2* and *Ncam1*, respectively.

Given the key role played by hydrogen bond interactions, we decided to analyze the dynamics of hydrogen bond networks for the trajectories of *Adamts14*-*1* and *Ncam1*-*ext* in more detail. Interestingly, the hydrogen bonding dynamics are markedly different between *Adamts14*-*1* and *Ncam1*-*ext*, despite the very similar RNA-DNA interface (i.e., RNA-pyrimidine/DNA-purine pairs, pyr/d-pur, only interrupted by one C-dT mismatch and one G-dG mismatch, immediately followed by an A-dT pair). Specifically, we first evaluated the frequency of the so-called “*in-register*” hydrogen bonds, i.e., hydrogen bonds that are formed between adjacent RNA and DNA bases when the strands are aligned. For *Adamts14*-*1*, *in-register* H-bonds are almost absent +/-four base pair steps around the A-dT/G-dG double pair (Figure 4d), whereas in *Ncam1*-*ext* an almost intact in-register hydrogen bond network can be found in the 5’-direction (Figure 4e). Next, out-of-register hydrogen bonding across the RNA and the DNA strands was also evaluated. Both *Adamts14*-*1* and *Ncam1*-*ext* display similar and highly dynamic patterns of out-of-register hydrogen bonds, forming transiently and not simultaneously in different parts of the system, and with lower frequency with respect to the in-register ones (Supplementary Figure S8a and b).

These analyses suggest that the *Ncam1* sequence has a higher probability to form a sequence-specific triple helix with *Snhg15* compared to *Adamts14*, due to its ability to establish a network of in-register hydrogen bonds (Figure 4). Thus, molecular modeling supports the experimental hypothesis that *Snhg15* forms a sequence-specific triple helix with the *Ncam1* locus, enabling the recruitment of DNMT1 to this genomic site, which can be modulated by ephrinA5-Fc triggered signaling. Moreover, these findings are reflected by our molecular biology analysis which detected changes in the methylation status for *Ncam1*, but not *Adamts14* (Supplementary Table S3), and only showed DNMT1 association in the promoter region of *Ncam1*.

## Discussion

We here provide evidence, that ephrinA5 acts on cell motility through *Snhg15-*mediated transcriptional regulation by orchestrating DNMT1 recruitment to the *Ncam1* locus thereby facilitating DNA methylation-dependent repression. As *Snhg15* is a cancer-related lncRNA, and CB cells serve as a model for medulloblastoma (Behesti & Marino, 2009; Gilbertson & Ellison, 2008), these findings might have impact on tumor cell biology, since the Eph/ephrin system is critically involved in cancer trajectories (Anderton et al., 2021).

Signals from the local microenvironment of a cell influence diverse aspects of cell intrinsic processes, key for tissue and organ development, whereas perturbances can lead to the initiation and progression of cancer (Casal, 2002; Herceg & Vaissière, 2011; Parsa, 2012). Typically, these signals are detected by membrane receptors, such as the Eph receptor tyrosine kinases, expressed in diverse tissues including the mammalian brain, where they orchestrate neurodevelopmental processes through the regulation of critical aspects relevant for cell proliferation, differentiation, adhesion, and neuronal migration (Gerstmann et al., 2015; Gerstmann & Zimmer, 2018; Rudolph et al., 2010; 2014; Zimmer et al., 2007; 2008; 2011). Furthermore, the expression of Eph receptors and ephrins as well as dysregulations in their bidirectional signaling was suggested to play a crucial role in tumor formation (Pasquale, 2010), e.g., in glioma (Ferluga & Debinski, 2014) and medulloblastoma (Anderton et al., 2021; Bhatia et al., 2015; Sikkema et al., 2012). Ephrin-related signaling can act as both, a tumor suppressor and tumorigenic, depending on the activated downstream pathways.

The regulation of cell motility and migration is a key function of the Eph/ephrin family during organ development (e.g., the brain) as well as in cancer. Apart from an Eph/ephrin signaling-dependent modulation of actin dynamics (Kindberg et al., 2021), Eph receptors were shown to act on migration by mediating cell adhesion to the ECM (Nakada et al., 2005; Pasquale, 2010) through the activation of Ras homolog family member A (RhoA) via Src or the focal adhesion kinase (FAK) (Wang et al., 2020). Here, we found an ephrinA5-dependent increase in *Ncam1* expression, which is implicated in establishing cell-ECM interactions and is known to curtail cellular motility (Blaheta et al., 2002; Edvardsen et al., 1994; Prag et al., 2002). In contrast to its effects on CB cells, ephrinA5 was reported to increase the motility of embryonic cortical neurons (Zimmer et al., 2007) and to act as a repellent cue for embryonic neurons deriving from the medial ganglionic eminence, the source of cortical inhibitory interneurons (Zimmer et al., 2008). This emphasizes cell-type-specific functions of ephrinA5, which could be mediated by distinct receptors and downstream signaling. Indeed, while in cortical and MGE cells EphA4 was described to be the interacting receptor (Zimmer et al., 2007; 2008), in CB cells it appears to be the EphA2 receptor which mediates the observed motility reducing effects. Similarly, in human hematopoietic stem and progenitor cells ephrinA5 was shown to enhance the migration by binding to EphA7 (Nguyen et al., 2017), while the same ligand negatively impacts the migration of primary hippocampal neurons in an EphA7-independent fashion (Meier et al., 2011), stressing the relevance of the cell type- and receptor-specific functions.

Physiological effects of Eph-receptor activation are further dependent on the activated downstream signaling pathways, which can be modulated by the interactions of Eph receptors with other cell surface receptors, such as the fibroblast growth factor receptor (FGFR) and chemokine receptors, as well as cell adhesion molecules such as ß-integrins (Arvanitis & Davy, 2008). This even can lead to ligand-independent receptor activation (Chastney et al., 2020; Fang et al., 2008; Finney et al., 2021). However, as a knockdown of EphA2 without a stimulation with ephrinA5-Fc caused no significant changes in the motility of CB cells (Supplementary Figure S3a and b), only the ligand-induced activation seems relevant in this context.

Due to their implication in cell motility regulation, it is not surprising that both ephrinA5 and EphA2 have been described to influence tumorigenesis and tumor progression (Ireton & Chen, 2005; Li et al., 2009; WalkerLJDaniels et al., 1999; Wang et al., 2012). EphrinA5 was described to act as a tumor suppressor in glioma by negatively regulating the epidermal growth factor receptor (EGFR) (Li et al., 2009). In line with this, H3K27me3-mediated repression of ephrinA5 was suggested to promote tumor growth and invasion in glioblastoma multiforme (GBM) (Ricci et al., 2020). Likewise, EphA2 has been proposed as a tumor suppressor, which is upregulated at transcript and protein levels in human tissue samples and cancer cell lines (Miao & Wang, 2012; Pasquale, 2010; Tandon et al., 2011). However, EPHA2 can also act as an oncogenic protein, promoting migration, e.g., by ligand-independent activation of EPHA2 via Akt, whereas ligand-dependent activation was shown to abolish the promotion of cell motility (Miao et al., 2009), with the latter being in line with our findings.

Ephrin-triggered Eph-receptor activation has been reported to converge on pathways that signal to the nucleus, such as the MAPK/ERK and PI3K-Akt/PKB pathway (Arvanitis & Davy, 2008; Liang et al., 2019). Hence, besides remodeling focal adhesive complexes and the cytoskeleton, physiological responses (here: cell motility) could rely on induced transcriptional changes involving these pathways. Yet, so far, Eph/ephrin signaling triggered alterations in gene expression as well as the related gene regulatory mechanisms in the nucleus, are still under-investigated. Microarray-based analyses of cortical tissue from ephrinA5-deficient mice revealed essential and biologically significant transcriptional alterations (Peuckert et al., 2008). Further evidence for ephrinA5-dependent modulation of gene expression was provided by Meier et al. (2011), who reported an ephrinA5-mediated suppression of the BDNF-evoked neuronal immediate early gene response. Another ligand, ephrinA1, regulates hepatoma cell growth by triggering transcriptional changes of associated genes *in vitro* (Iida et al., 2005). In line with this, we identified changes in ECM- and migration-related gene expression in CB cells after 24 h of ephrinA5-Fc stimulation in a previous study (Pensold et al., 2021). Here, we report that about half of the differentially expressed genes also display changes in DNA methylation signatures, among which cell adhesion-related genes were significantly enriched. The gene coding for NCAM1, a neuronal cell adhesion molecule with key features for motility regulation in neurons as well as in cancer cells (Cui et al., 2019; Guan et al., 2020; Maness & Schachner, 2007; Schmid & Maness, 2008), was significantly increased in expression and its promoter region showed a significant reduction in CpG methylation after ephrinA5 stimulation. Since abolishing the ephrinA5-Fc-induced increase in *Ncam1* transcript levels rescued the motility impairments, ephrinA5-triggered transcriptional changes of cell adhesion-related genes seem to be implicated mechanistically in mediating this physiological response.

A negative correlation of *Ncam1* expression and cell motility has been described in the context of cancer, for instance for ameloblastoma cells and neuroblastoma (Blaheta et al., 2002; Guan et al., 2020). Moreover, *Ncam1* expression correlates positively with the survival rate in low-grade glioma patients (Supplementary Figure S2a). NCAM1 is a well-known tumor suppressor in numerous cancer types (Katoh & Katoh, 2003; Roesler et al., 1997). In line with our findings, *NCAM1*-transfected glioma cells were reported to migrate less (Prag et al., 2002) and they exhibit decreased invasiveness upon transplantation into the rat brain (Edvardsen et al., 1994).

Since Eph/ephrin signaling is fundamentally involved in tumorigenesis as well, ephrinA5-triggered Eph-signaling could be a potential pathway that leads to diminished *NCAM1* expression in cancer cells. As DNA methylation signatures of *Ncam1* and numerous other genes were essentially remodeled after ephrinA5-Fc stimulation in CB cells, an Eph/ephrin dependent modulation of the DNA methylome may be a feasible mechanism of transcriptional regulation underlying reduced cellular motility. In fact, differential promoter methylation of genes encoding for adhesion molecules, such as the epithelial cell adhesion molecule (Ep-CAM) and E-cadherin, have been frequently linked to cancer cell motility, and invasion and metastasis of cancer (Chan et al., 2003; Chen et al., 2010; Tai et al., 2007).

LncRNAs were described to interact with proteins and DNA, e.g., via formation of triplex structures (Blank-Giwojna et al., 2019; Leisegang et al., 2022; Li et al., 2016), through which they could potentially mediate locus-specific epigenetic remodeling by recruiting or evicting epigenetic modifiers to discrete DNA loci (Zimmer-Bensch, 2019). These remodeling processes have been implicated in cell physiological functions, including cell proliferation, differentiation, and migration (Mercer et al., 2009). Importantly, changes in lncRNA expression can lead to transcriptional dysregulation of diverse gene sets. Such aberrant expression patterns of lncRNAs have been observed in the pathophysiology of various diseases including cancer, where they are proposed to play role in metastasis and the remodeling of the tumor environment (Fang & Fullwood, 2016; Jiang et al., 2019).

We also found dysregulated expression of lncRNAs such as *Snhg15*, an important cancer-related lncRNA, in response to ephrinA5-Fc treatment. Its human orthologue *SNHG15,* which we identified to be similarly diminished in expression after ephrinA5-Fc stimulation (Pensold et al., 2021), has been reported to be upregulated in multiple types of cancer. *SNHG15* participates in the initiation and progression of diverse cancer types by affecting proliferation and migration (Tong et al., 2019). The pro-oncogenic and pro-migratory function of *SNHG15* is in line with the ephrinA5-Fc-induced downregulation of *Snhg15* in CB cells, which are commonly used as a medulloblastoma cell model (Behesti & Marino, 2009; Pensold et al., 2021), as well as the observed motility restriction. *SNHG15* has been often demonstrated to have a sponging function, binding and disabling various miRNAs to upregulate the expression of oncogenic genes in glioma, breast cancer, and lung cancer (Jin et al., 2018; Kong & Qiu, 2018; Y. Ma et al., 2017). Yet, RNA immunoprecipitation (RIP) assays have revealed that *SNHG15* can also interact with EZH2 to repress tumor suppressor genes via EZH2-mediated trimethylation at H3K27 in the nucleus (Z. Ma et al., 2017). While no such interaction between *Snhg15* and EZH2 was detected in CB cells by CLIP, which in contrast to RIP captures direct RNA-protein interactions, we found an interaction of *Snhg15* with DNMT1, a major DNA methyltransferase (Bestor, 2000; Mohan & Chaillet, 2013; Pensold & Zimmer-Bensch, 2021; Svedruzic, 2008; 2011). Furthermore, this interaction was reduced upon ephrinA5-Fc stimulation. DNMT1 has already been reported to interact with lncRNAs, e.g., in colon cancer, and the deregulation of DNMT1-associated lncRNAs was proposed to contribute to aberrant DNA methylation and gene expression in colon tumorigenesis (Merry et al., 2015). Another lncRNA, *NEAT1*, interacts with DNMT1, orchestrating cytotoxic T-cell infiltration in lung cancer (Ma et al., 2020). There is increasing evidence that apart from pathophysiological conditions, DNMT-lncRNA interactions and lncRNA-mediated DNA methylation are likewise important for normal cell physiological regulation (Huang et al., 2022).

In line with this, we here provide evidence for *Snhg15-*dependent recruitment of DNMT1 to and DNA methylation of the *Ncam1* promoter region, which was abolished after ephrinA5-Fc treatment. Using sequence-based algorithms, we characterized putative DNA binding sites of *Snhg15* at promoter regions of *Ncam1* and *Adamts-14.* Sequence based algorithms, which consider canonical base pairing rules driving RNA-DNA triple helices (Buske et al., 2011; Warwick et al., 2023) to find potential triple helices, are commonly used in the literature. These fast algorithms can evaluate and rank thousands of RNAs and putative DNA target sequences, which commonly arise in sequencing experiments (Kuo et al., 2019). This allowed us to characterize the potential role of *Snhg15* in triple helix mediated DNA interactions (Pensold et al., 2021). These sequence-based methods provide hypotheses on the positions, size and alignments of the interaction patterns that are parameters necessary to build targeted all atom models, which then provide further insight into the physics of the interaction, such as stability, dynamics, and role of individual nucleotides.

Therefore, next atomistic models were constructed based on the predicted binding site alignments. Specifically, atomistic model not only helped in selecting the most stable triple helix among the ones predicted for both *Adamts14* and *Ncam1,* but also, their ability to establish specific interactions with *Snhg15* was evaluated. Molecular dynamic simulations suggest that while the selected triple helices for *Ncam1* and *Adamts14* display a comparable overall stability, the local interaction established at the RNA-DNA interface is significantly different. *Ncam1* indeed features several *in-register* hydrogen bonds, persistent over the entire simulations time, that maintain the sequence-specific complementary of hydrogen bond networks between the RNA-DNA triplex interfaces. This is not the case for *Adamts14* where the lack of persistent *in-register* hydrogen bonds suggests a less sequence-specific interaction between RNA and DNA. Such results are in line with our wet lab approaches: for the *Ncam1* but not for *Adamts14* promoter region significant change in methylation levels were detected in response to ephrinA5-Fc treatment.

There are only a few simulations of RNA/DNA interaction via triple helix formation (Kunkler et al., 2019; Leisegang et al., 2022). Optimizing model details and parameters for such molecular dynamics simulations, such as the protonation state and appropriate force fields, remains an active area of research (Antonov et al., 2019) which is yet still hampered by the so-far low number of physical experiments on these molecules.

The lncRNA-mediated targeting of epigenetic writers such as DNMTs or histone modifying proteins represents an attractive mechanism for dysregulation of epigenetic signatures that occurs in cancer cells. This finding seems to be specifically relevant since the regulation of lncRNA expression is responsive to signaling from peripheral membrane receptors commonly reported in cancer. While we identified Eph/ephrin signaling to modulate lncRNA expression (Pensold et al., 2021), altered neuronal activity, depending on ion channel opening in the cell membrane, also leads to changes in lncRNA expression (Barry et al., 2014; Barry et al., 2017; Lipovich et al., 2012). As glioma progression is modulated by both the Eph/ephrin signaling and neuronal activity, this could lead to specific alterations in the lncRNA repertoire which in turn act on transcription and translation at different levels. Similar mechanisms are conceivable for medulloblastoma and other types of cancer. However, how (other) distinct lncRNAs are affected in their transcription in response to, e.g., ephrin stimulation or neuronal activity, and which signaling pathways are involved, needs to be dissected in future studies.

## Data Availability

The Illumina Mouse Methylation BeadChIP data have been deposited with GEO-NCBI, the accession number GSE will be provided latest upon acceptance of the manuscript (in progress).

## Acknowledgements

This work was supported by the Deutsche Forschungsgemeinschaft (DFG, German Research Foundation) [368482240/GRK2416 to GZB and GR, ZI 1224/8-1 to GZB, ZI 1224/13-1 to GZB]; the Excellence Initiative of the German federal and state governments, RWTH Aachen, Seed Fund, grant number OPSF678 to GR, OZ, IC and GZB. The graphical abstract was created with BioRender.com.

## Figure legends

**Supplementary Figure S1: CLIP revealed no interaction of *Snhg15* and EZH2 in CB cells.** RNA recovery for IgG and anti-EZH2 antibody CLIP samples in CB cells (N = 5 biological replicates, Tukey und Spear box plot). For all investigated amplicons, the recovery for EZH2 could not be statistically differentiated from the IgG-based pulldown. Significances were determined with two-tailed Student’s *t*-test. ctrl-Fc: control-Fc, efnA5-Fc: ephrinA5-Fc, CB: cerebellar granule, CLIP: UV cross-linking and immunoprecipitation.

**Supplementary Figure S2: NCAM1 is implicated in the survival rate of low-grade glioma patients and its expression can be downregulated in murine cells via RNA silencing.** (a) High expression levels of *NCAM1* are associated with increased patient survival in low-grade glioma. Survival analysis is based on clinical data and gene expression counts from tumor samples of lower grade glioma patients downloaded from BioPortal (http://www.cbioportal.org/study/clinicalData?id=lgg_tcga) and The Cancer Genome Atlas (TCGA), respectively. (b) Knockdown efficiency of the applied *Ncam1* siRNA (N = 3 biological replicates). Significances were determined with log-rank (a) and Wilcoxon-Mann-Whitney test (b). Significance levels: *p* value < 0.05 *; *p* value < 0.01 **; *p* value < 0.001 ***. ctrl: control. LGG: low-grade glioma. siR: siRNA.

**Supplementary Figure S3: Migratory analysis of CB cells upon stimulation with ephrinA5-Fc and downregulation of *EphA2*.** The motility of CB cells was reduced upon stimulation with ephrinA5-Fc. (a) Temporal color-coded migratory distance over 20 h of imaging. The starting point of migration for each cell is shown in dark blue and the end point in white. (b) Quantitative analysis of average migratory speed (n = 268 for ctrl siR + ctrl-Fc, n = 244 for ctrl siR + efnA5-Fc, n = 273 for *EphA2* siR + ctrl-Fc, n = 238 for *EphA2* siR + efnA5-Fc, N = 3 biological replicates). (c) Knockdown efficiency of the applied *EphA2*-siRNA (N = 3 biological replicates). Significances were determined with one-way ANOVA (b) and Wilcoxon-Mann-Whitney test (c). Significance levels: *p* value < 0.05 *; *p* value < 0.01 **; *p* value < 0.001 ***. Scale bar: 100 μm. ctrl: control. efnA5: ephrinA5. siR: siRNA.

**Supplementary Figure S4:** Native ChIP reveals no changes in DNMT1 association and histone methylation signatures within the proximity of a *cis*-regulatory element of *Adamts14* and the putative *Snhg15* binding sites. (a-c) ChIP-qPCR analysis with anti-DNMT1, anti H3K27me3 and anti-H3K4me3 antibodies for the promoter region of *Adamts14* normalized against the input material and IgG (N = 3 biological replicates for *P1* (a), N = 4 biological replicates for *P2* (b-c)). (d) Genomic regions targeted by the primers are shown in turquoise. The promoter is shown in dark blue, candidate cis-regulatory elements in red, and putative *Snhg15* binding sites in pink. Significances were determined with two-tailed Student’s *t*-test. Significance levels: *p* value < 0.05 *; *p* value < 0.01 **; *p* value < 0.001 ***. ctrl: control. efnA5: ephrinA5.

**Supplementary Figure S5:** Root mean square deviation (RMSD) plot for *Ncam1*, *Ncam1*-*ext*, *Adamts14-1* and *Adamts14*-*2* taken over the entire trajectory of 600 ns for (a) the RNA strand of the triple helix and (b) the DNA double helix.

**Supplementary Figure S6:** Comparison of energy contributions between the two *Adamts14* models (*Adamts14*-*1* and *Adamts14*-*2*). The hydrogen bond energies between individual residues at the same base pair level (i) were calculated by taking the sum of Lennard-Jones (LJ) and Coulomb (CB) short range interaction energies. The cross energies were calculated for a base pair level (i) by considering the sum of LJ and CB short range interaction energies of (i)th residue in chain B with (i+1)th and (i-1)th residue in chain A and C. The stacking energies were calculated for a base pair step level by considering the sum of LJ and CB short range interaction energies of (i)th and (i+1)th residues in chain A, B and C. The total plot shows the sum of the contribution of individual energies at the (i)th base pair level. In all calculations the energy contributions involving terminal residues were not included to avoid discrepancy in the number of terms that contribute.

**Supplementary Figure S7:** Comparison of energy contributions between the two *Ncam1* models (*Ncam1-1* and *Ncam1*-*ext*). The hydrogen bond energies between individual residues at the same base pair level (i) were calculated by taking the sum of Lennard-Jones (LJ) and Coulomb (CB) short range interaction energies. The cross energies were calculated for a base pair level (i) by considering the sum of LJ and CB short range interaction energies of (i)th residue in chain B with (i+1)th and (i-1)th residue in chain A and C. The stacking energies were calculated for a base pair step level by considering the sum of LJ and CB short range interaction energies of the (i)th and (i+1)th residue in chain A, B and C. The total plot shows the sum of the contribution of individual energies at the (i)th base pair level. In all calculations the energy contributions involving terminal residues were not included to avoid discrepancy in the number of terms that contribute.

**Supplementary Figure S8:** Display of all hydrogen bonded interactions of the RNA that occur with frequency >1% in the (a) *Adamts14*-*1* and (b) *Ncam1*-*ext* simulation. For clarity the purine DNA strand was drawn above, and the pyrimidine DNA strand below the RNA. H-bonds between the two DNA strands as well as within individual strands are omitted to focus on the H-bond patterns of the RNA strand. Occurrence is in “H-bond units”, i.e., an occurrence of >100% indicates that on average there exists more than one H-bond between the respective residues.

**Supplementary Table S1:** Sequences of applied primer pairs in (RT-)qPCR experiments.

**Supplementary Table S2:** Detailed sequences of the simulated systems (*Adamts14*-*1*, *Adamts14*-2*, Ncam1, Ncam1*-ext). Pairing between DNA sequences (black) and RNA sequences (red) is predicted to form triple helixes for two binding sites in the *Adamts14* promoter and one binding site at the *Ncam1* promoter. 5’ and 3’ indicates the orientation of the DNA and RNA strands. “|” indicates base pairing following triple helix canonical code, while “*” indicates positions with a mismatch.

**Supplementary Table S3.** Differentially in CB cells treated with ephrinA5-Fc. Table lists all probes/regions (DMRs) with an adjusted *p* value ≤ 0.05 (adj. *p*val).

**Supplementary Table S4:** Table depicts all 19 protein-coding genes upregulated after 24 h ephrinA5-Fc treatment and with putative triplex target DNA sites (TTS) for *Snhg15* (RNA-seq; Pensold et al. 2021).

## Supplementary Methods

### Survival Analysis

The survival analysis was implemented in R. It was based on clinical data and gene expression counts from tumor samples of lower grade glioma patients downloaded from BioPortal (http://www.cbioportal.org/study/clinicalData?id=lgg_tcga) and The Cancer Genome Atlas (TCGA), respectively. To download the gene expression counts, we applied the R package TCGAbiolinks setting the parameters for GDCquery as follows: project=”TCGA-LGG”, data.category=”Transcriptome Profiling”, data.type=”Gene Expression Quantification”, workflow.type=“HTSeq – Counts” and sample.type=c(“Primary solid Tumor”, “Recurrent Solid Tumor”). The raw gene counts were normalized by applying DESeq2::varianceStabilizingTransformation(). Afterwards, the patients were divided into two groups (high expression, low expression) based on whether their NCAM1 expression level is above or below the median NCAM1 expression level among all patients. Using survival::Surv() and survival::survfit() while setting the time parameter to the survived months and the event parameter to the patients’ vital status from the clinical data a Kaplan-Meier curve object was created. It was plotted with survminer:: ggsurvplot() setting risk.table=TRUE. If not indicated otherwise, default parameters have been passed to the applied R functions.

