## Supplementary material (Figures and Tables) for "EphrinA5 regulates cell motility by modulating the targeting of DNMT1 to the *Ncam1* promoter via lncRNA/DNA triplex formation"

### Supplementary Figure 1

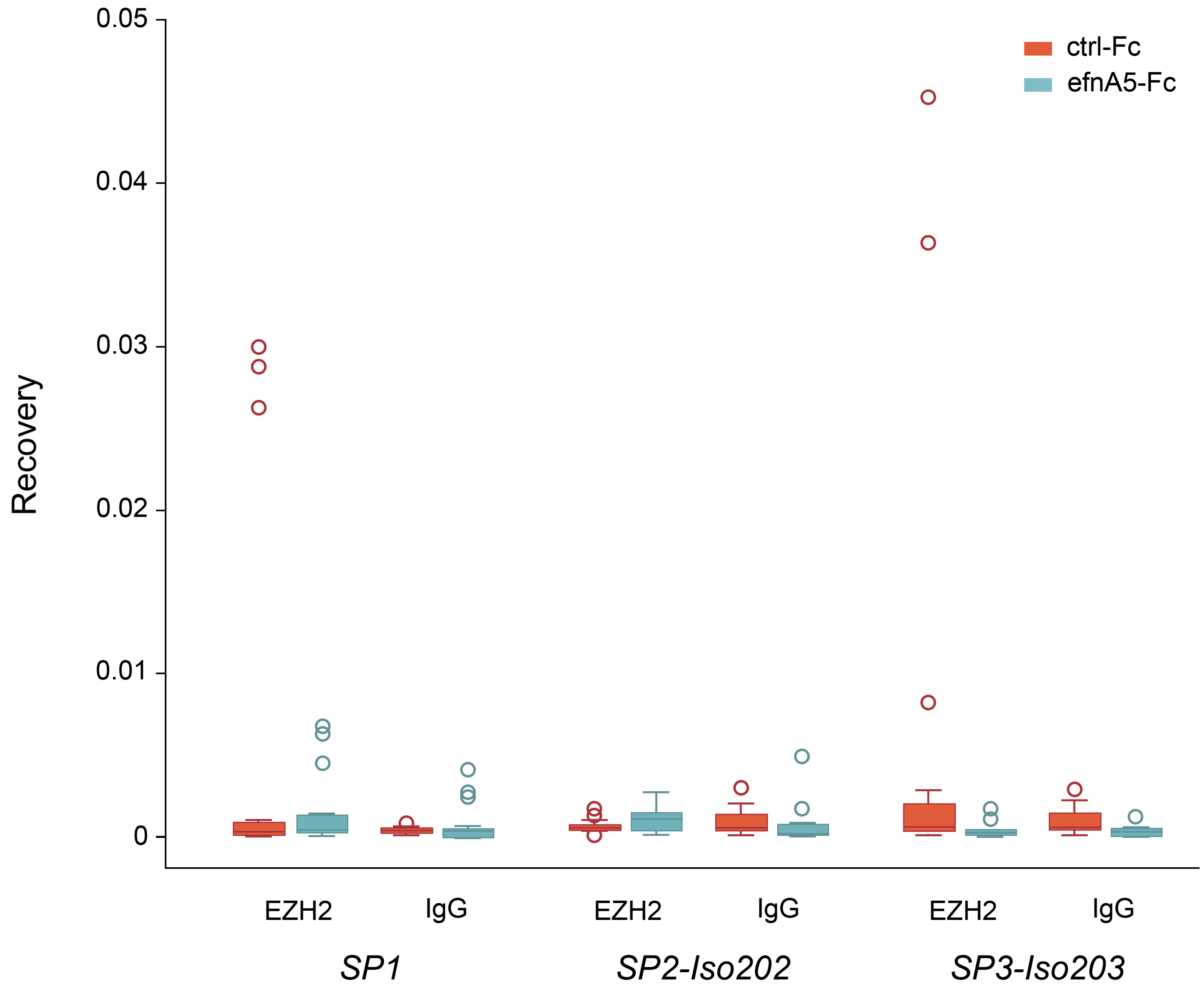

### Supplementary Figure 2:

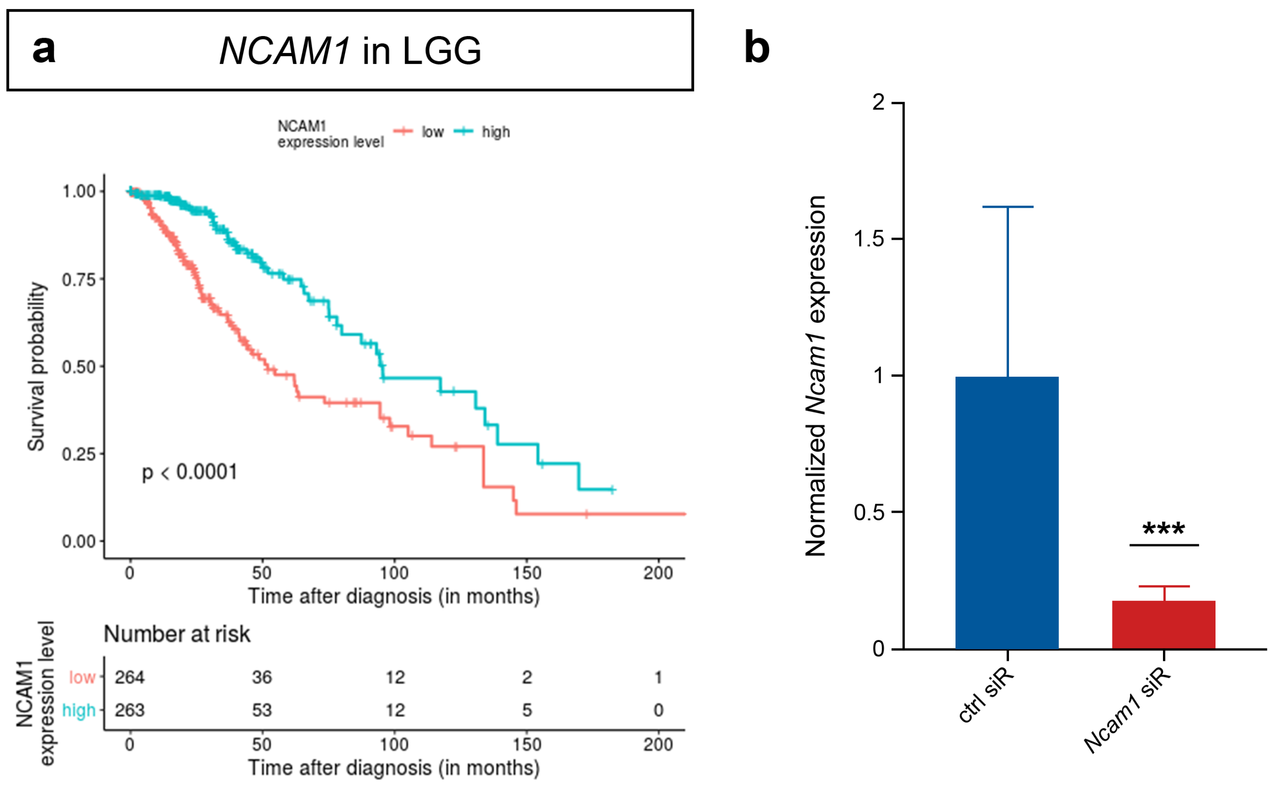

### Supplementary Figure 3:

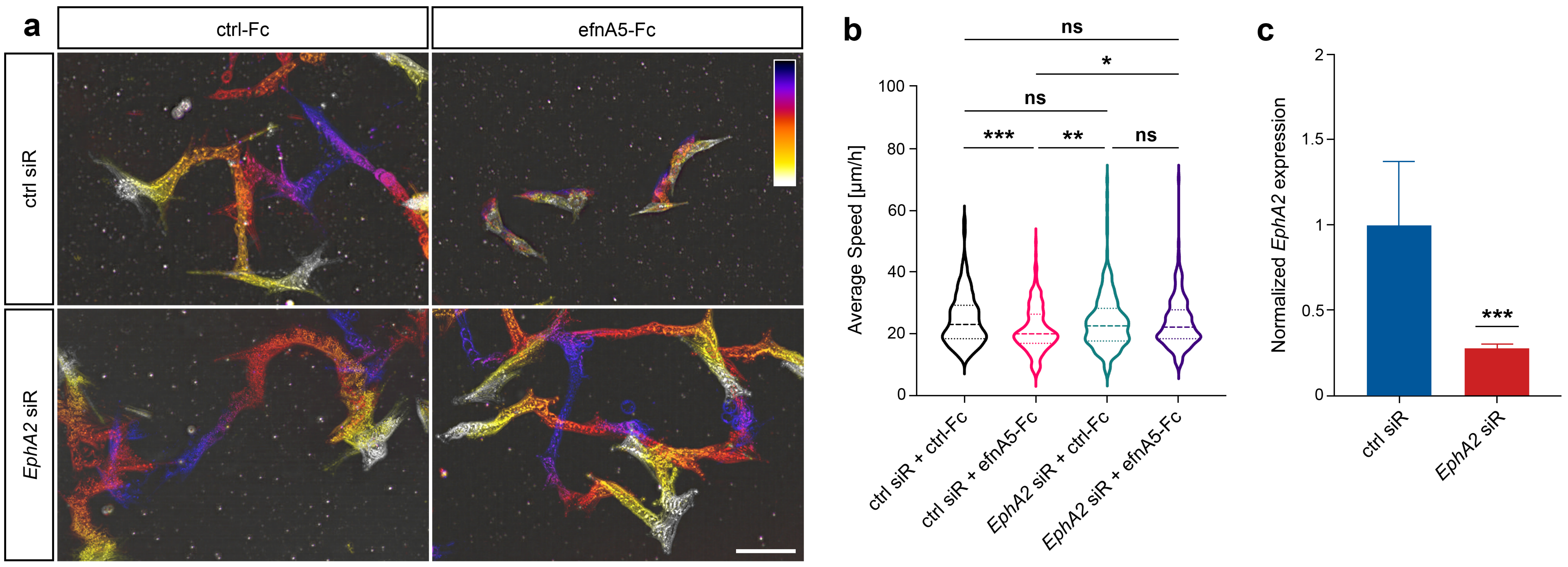

### Supplementary Figure S4:

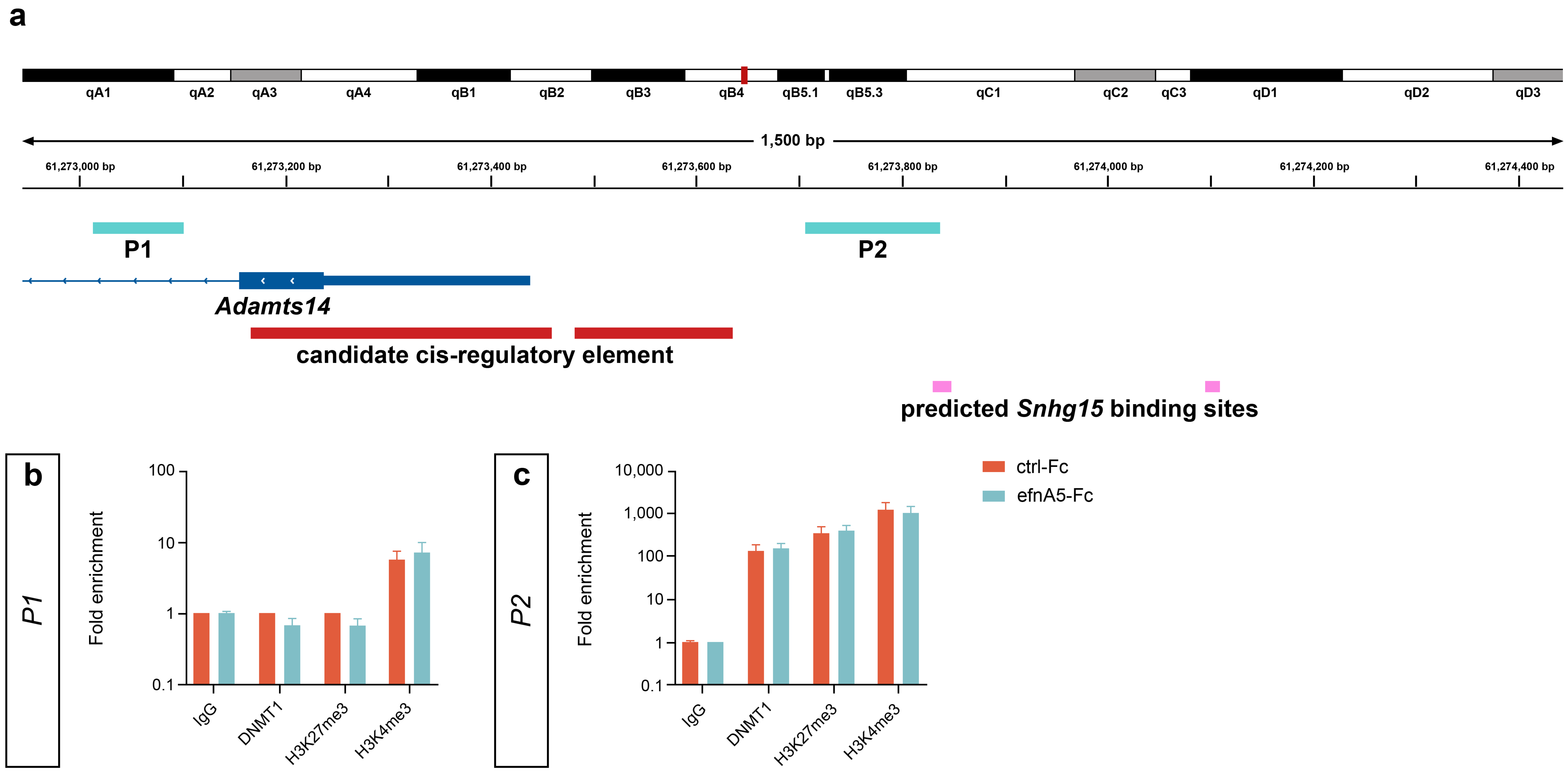

### Supplementary Figure S5:

**a**

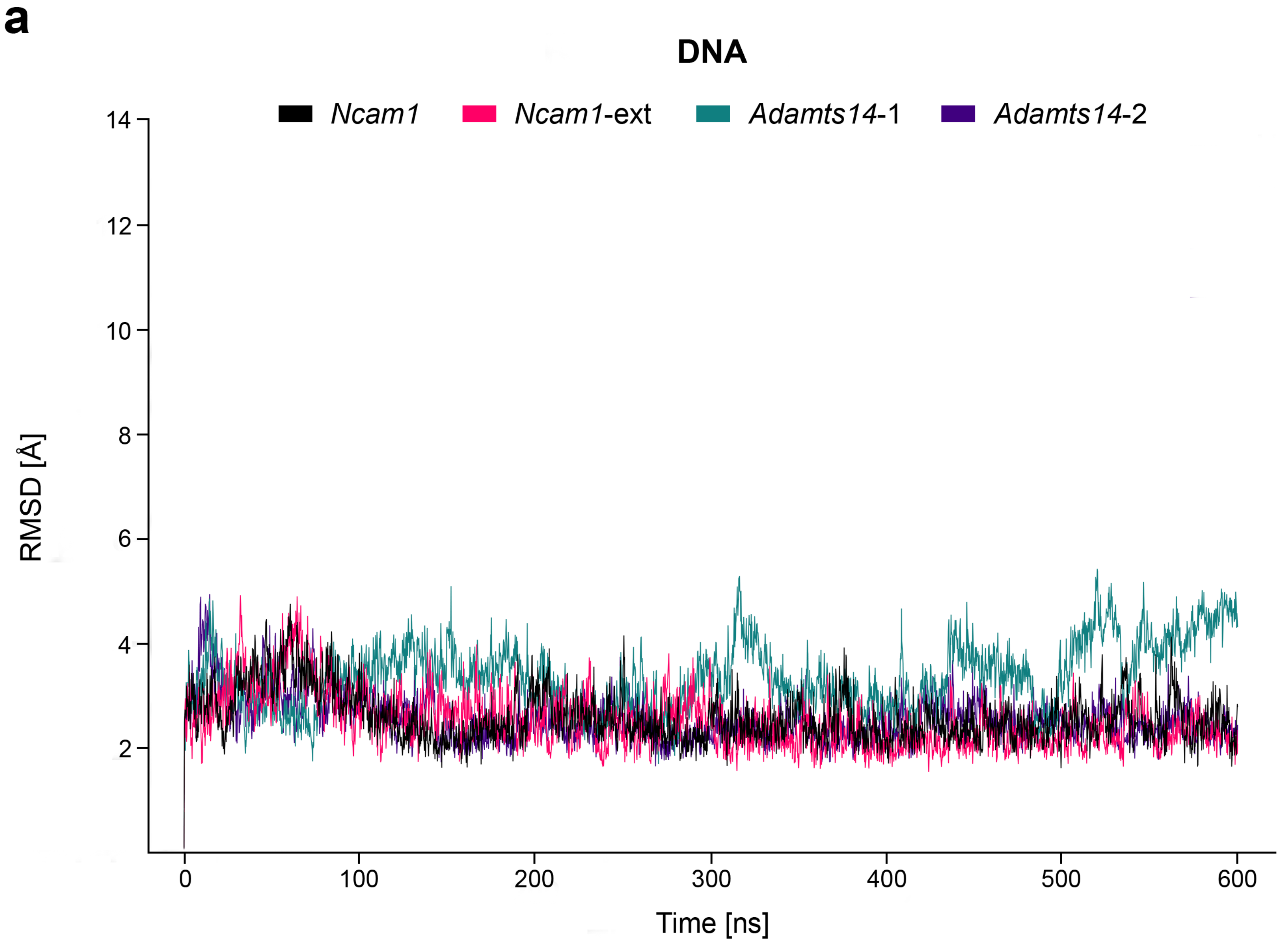


**b**

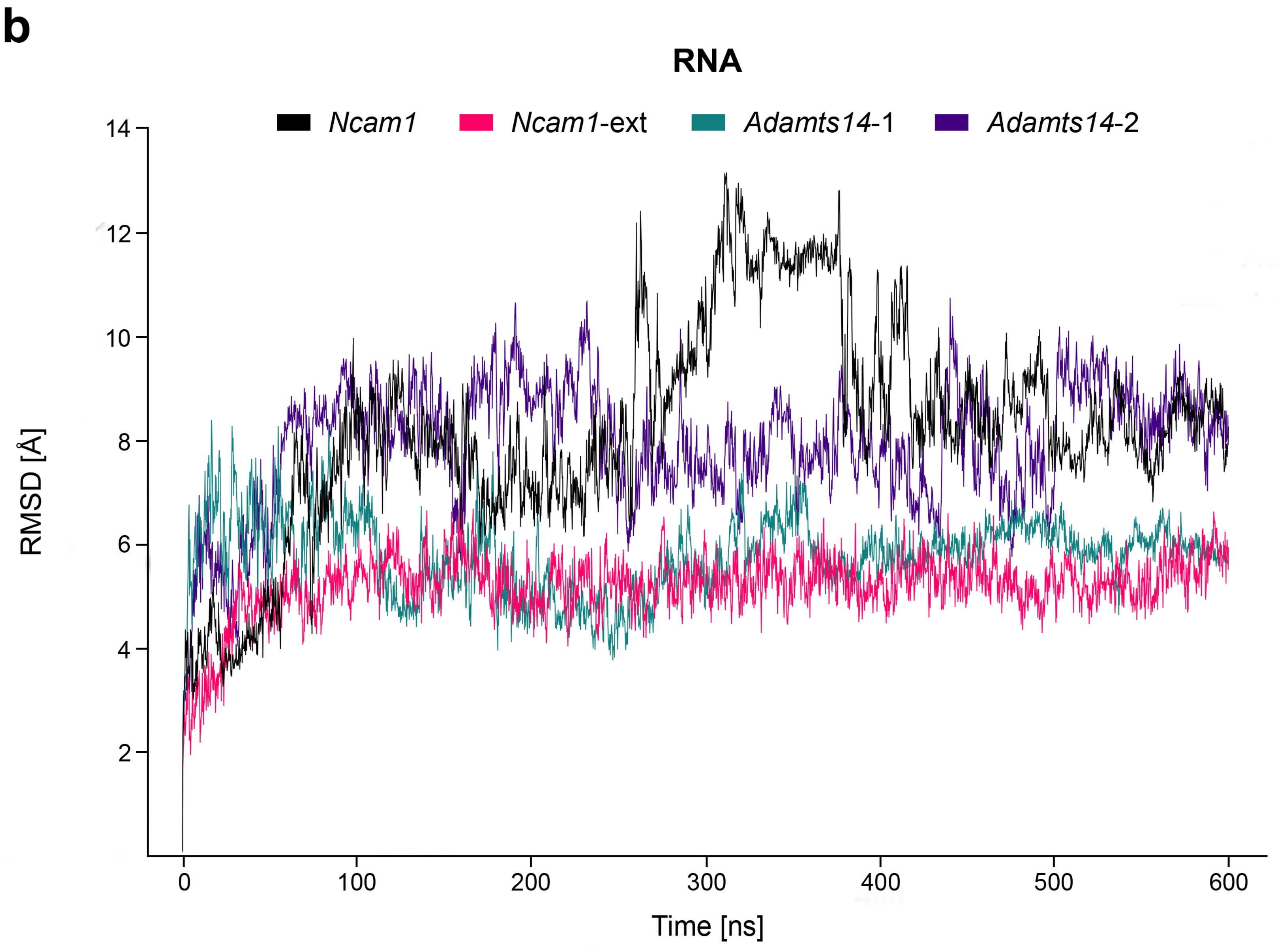

### Supplementary Figure S6:

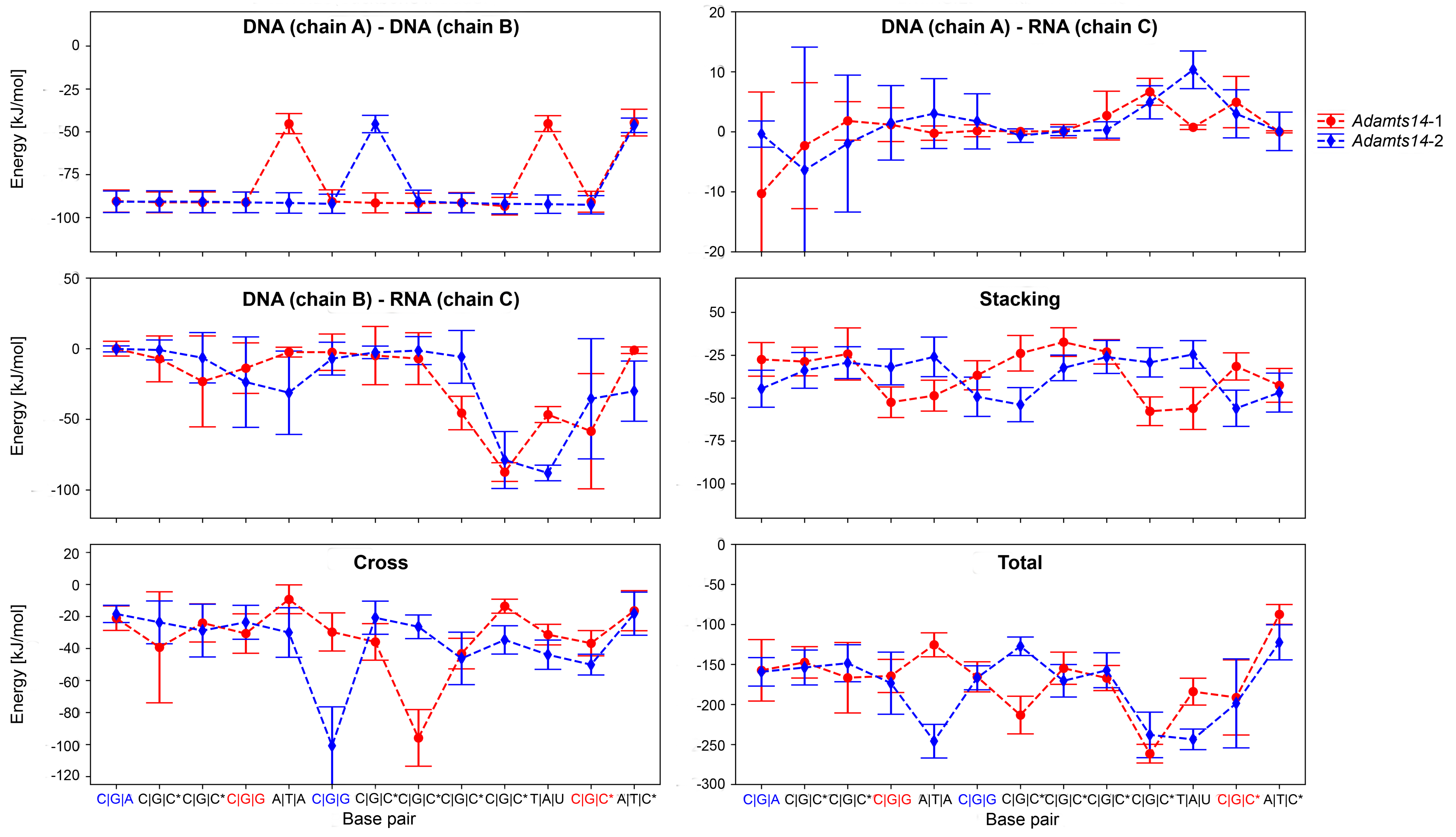

### Supplementary Figure S7:

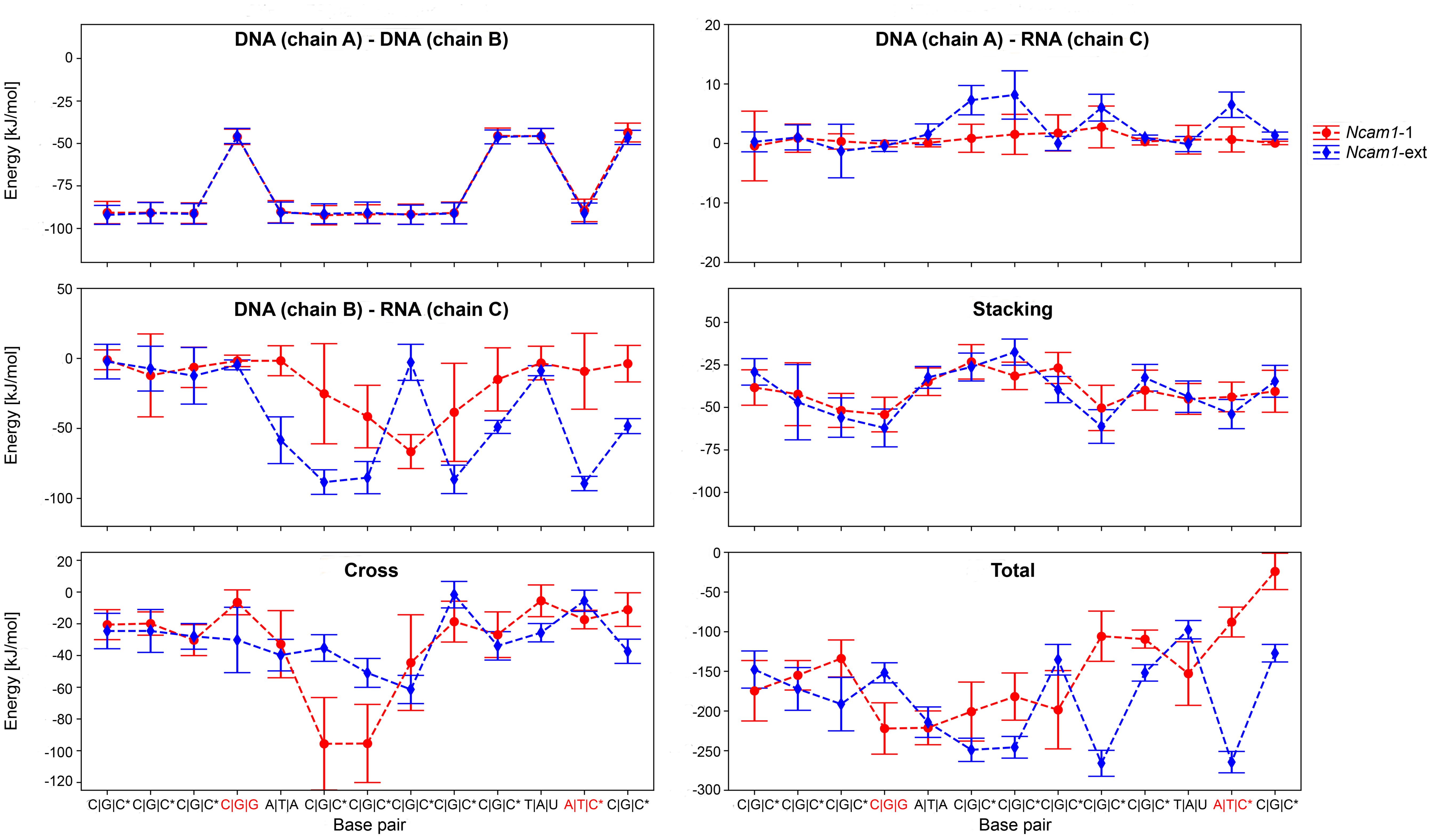

### Supplementary Figure S8:

**a** *Adamts14*-1

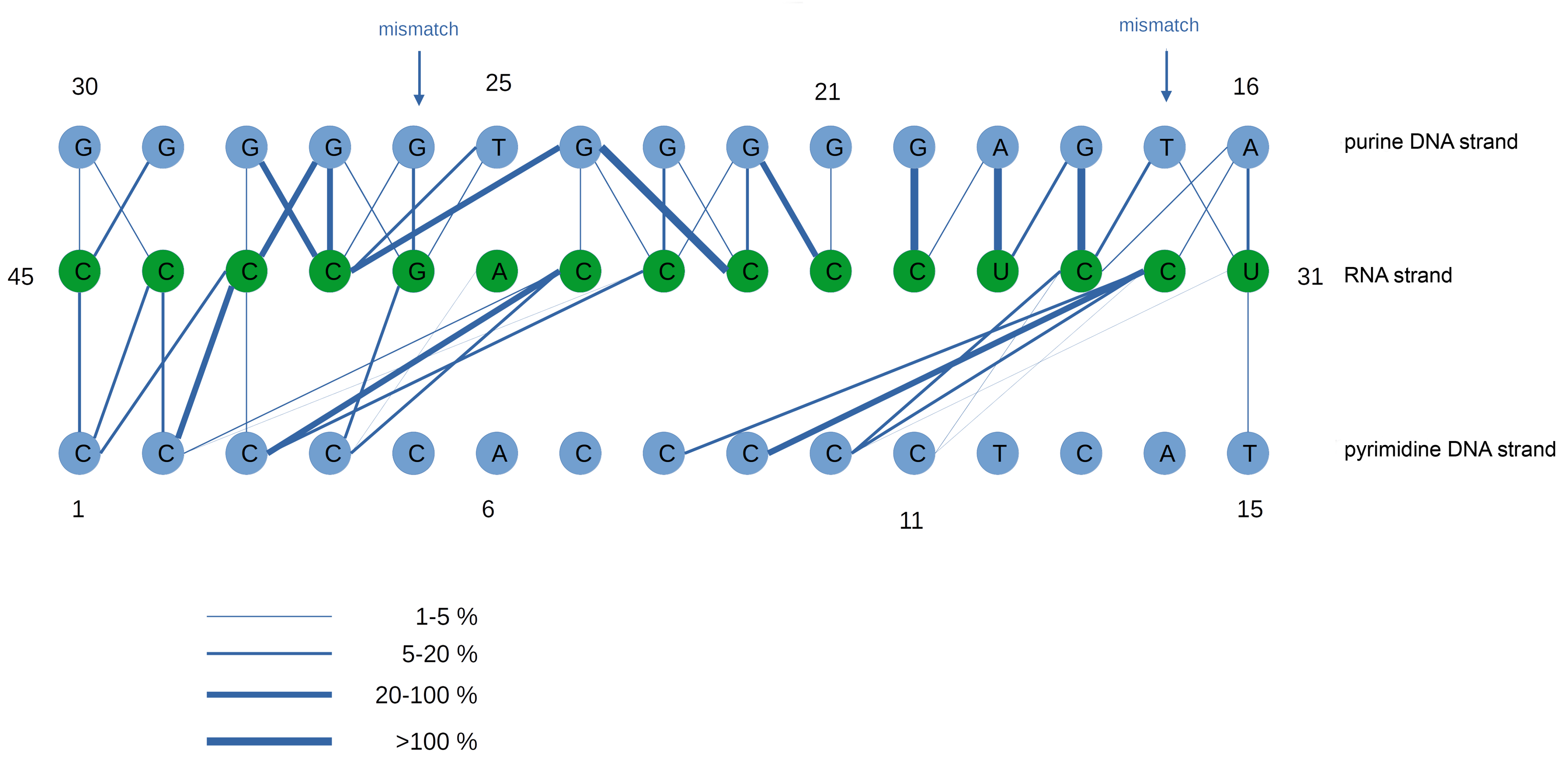


**b** *Ncam1*-ext

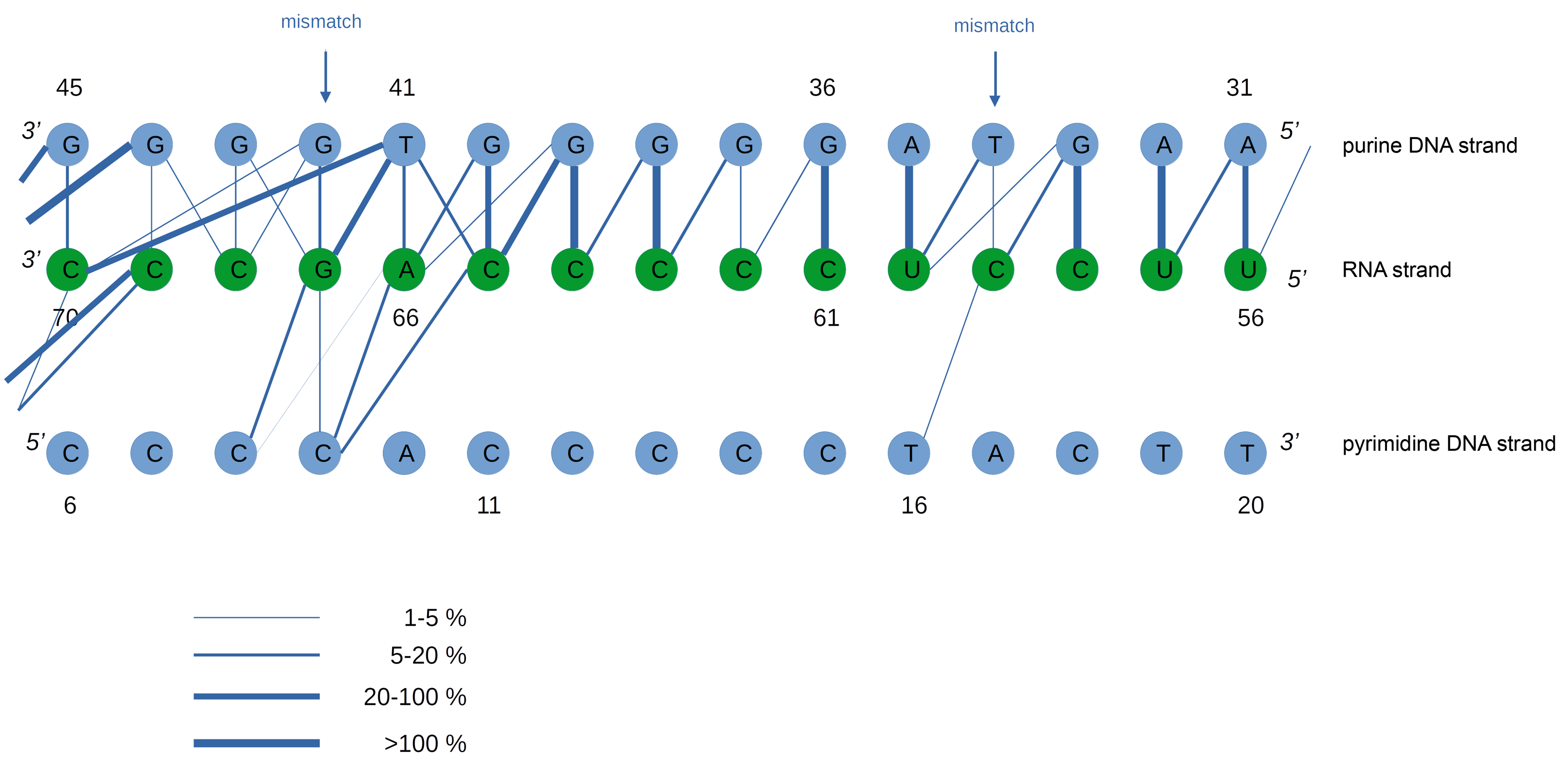

### Supplementary Table S1:

| Target | Descriptor | Sequence | Product size [bp] |
|---|---|---|---|
| *Adamts14* (DNA) | *P1* | F: AAAGCTAAGCAACTCAACCCAC<br>R: AGAGTGGCTCCTGGATGCAG | 90 |
| | *P2* | F: CATTGAATTTTGAAGGCGCAGATTG<br>R: CTCATATCGTTGTGACCTGACAG | 132 |
| *Atp5bp* (mRNA) | | F: TGTGATTACCCCAGAGACCT<br>R: TGCTCGACTGCTTTACTTCTTC | 149 |
| *EphA2* (mRNA) | | F: GTGTGCAAGGTGTCCGATTT<br>R: AACTTGCGGTAGGAAATGGC | 128 |
| *Ncam1* (DNA) | *P1* | F: TTATTCGGGGCGGGGAAGAC<br>R: CTAGAGGTGTTTGTCAAGCTTCGAG | 137 |
| | *P2* | F: GACAATTAACAGACCCGATGTAGAT<br>R: ACTAAGGATCTCATCTGGACTTTGT | 98 |
| | *P3* | F: TTTTCCTTCATTCACGTCTTTGGAG<br>R: GTGTCCACAACTTTGAAAATCGAAC | 129 |
| *Ncam1* (mRNA) | | F: GGTGACCCCTGATTCAGAAA<br>R: GGATGGAGAAGACGGTGTGT | 118 |
| *Snhg15* (lncRNA) | *SP1* | F: GCCCTCACCTTATCTTCCGT<br>R: AGCTCAGCCATCTCTACCTG | 78 |
| | *SP2* | F: AGCCTGTGTTCTTTTCTGGAG<br>R: AGGAAGAGTCAGTGAGCCAA | 145 |
| | *SP3* | F: CTACAGCGTGGAAGGGATGA<br>R: TCCCAGTCCCCAGAAGTCTA | 143 |

### Supplementary Table S2:

| Name (PDB code) | Description | Sequence |
|---|---|---|
| *Adamts14*-1 | DNA - DNA \| RNA | 5' C C C C C A C C C C C T C A T 3'<br>3' G G G G G T G G G G G A G T A 5'<br>\| \| \| \| * \| \| \| \| \| \| \| \| * \| \|<br>3' C C C C G A C C C C C U C C U 5' |
| *Adamts14*-2 | DNA - DNA \| RNA | 5' C C C C C C C A C C C C C T C 3'<br>3' G G G G G G G T G G G G G A G 5'<br>\| * \| \| \| \| * \| \| \| \| \| \| \| \|<br>3' C A C C C C G A C C C C C U C 5' |
| *Ncam1* | DNA - DNA \| RNA | 5' C C C C A C C C C C T A C T T 3'<br>3' G G G G T G G G G G A T G A A 5'<br>\| \| \| * \| \| \| \| \| \| \| * \| \| \|<br>3' C C C G A C C C C C U C C U U 5' |
| *Ncam1*-ext | DNA - DNA \| RNA | 5' G G C T T C C C C A C C C C C T A C T T C T G A G 3'<br>3' C C G A A G G G G T G G G G G A T G A A G A C T C 5'<br>\| * \| * * \| \| \| * \| \| \| \| \| \| \| * \| \| \| * * * * \|<br>3' C U C A C C C C G A C C C C C U C C U U A A C C G 5' |

### Supplementary Table S4:

| Promoter | Gene | TTSs Count | TTS Coverage |
|---|---|---|---|
| chr4:53159894-53160895 | *Abca1* | 31 | 0.31 |
| chr17:43800350-43801351 | *Rcan2* | 43 | 0.11 |
| chr3:89214179-89215180 | *Thbs3* | 49 | 0.08 |
| chr9:62537043-62538044 | *Coro2b* | 68 | 0.07 |
| chr9:13296956-13297957 | *Maml2* | 15 | 0.12 |
| chr15:83088505-83089506 | *Sehrl* | 15 | 0.07 |
| chr3:52105075-52106076 | *Maml3* | 14 | 0.07 |
| chr18:79109390-79110391 | *Setbp1* | 8 | 0.07 |
| chr8:82862355-82863356 | *Rnf150* | 5 | 0.048 |
| chr4:130046839-130047840 | *Col16a1* | 8 | 0.042 |
| chr4:114405723-114406724 | *Trabd2b* | 5 | 0.044 |
| chr7:98488282-98489283 | *Lrrc32* | 3 | 0.045 |
| chr10:61273437-61274438 | *Adamts14* | 4 | 0.032 |
| chr18:44812181-44813182 | *Mcc* | 2 | 0.031 |
| chr19:31765032-31766033 | *Prkg1* | 2 | 0.031 |
| chr2:173276532-173277533 | *Pmepa1* | 1 | 0.015 |
| chr9:49798924-49799925 | *Ncam1* | 76 | 0.23 |
| chr10:8518824-8519825 | *Ust* | 46 | 0.15 |
| chr18:33464028-33465029 | *Nrep* | 83 | 0.09 |
